# An angiosperm-wide perspective on reproductive strategies and floral traits

**DOI:** 10.1101/2024.02.26.582019

**Authors:** Andrew J. Helmstetter, Marcos Méndez, Jürg Schönenberger, Concetta Burgarella, Bruce Anderson, Maria von Balthazar, Sylvain Billiard, Hugo de Boer, Johanne Cros, Pierre-Alexandre Delecroix, Mathilde Dufay, John R. Pannell, Dessislava Savova Bianchi, Daniel J. Schoen, Mario Vallejo-Marin, Rosana Zenil-Ferguson, Hervé Sauquet, Sylvain Glémin, Jos Käfer

**Affiliations:** FRB-CESAB, 34000 Montpellier, France; ISEM, Univ Montpellier, CNRS, IRD, 34095 Montpellier, France; Area of Biodiversity and Conservation and Instituto de Investigación en Cambio Global (IICG), Universidad Rey Juan Carlos, 28933 Móstoles (Madrid), Spain; Department of Botany and Biodiversity Research, University of Vienna, Rennweg 14, 1030 Vienna, Austria; Department of Organismal Biology, University of Uppsala, 75236 Uppsala, Sweden; Department of Botany and Zoology, University of Stellenbosch, Private Bag X1, Matieland 7602, South Africa; University Lille, CNRS, UMR 8198 – Evo-Eco-Paleo, 59000 Lille, France; Natural History Museum, University of Oslo, 0318 Oslo, Norway; Université de Lyon, Université Lyon 1, CNRS, Laboratoire de Biométrie et Biologie Evolutive UMR 5558, 69622 Villeurbanne, France; CNRS, Ecosystèmes Biodiversité Evolution (Université de Rennes), 35000 Rennes, France; CEFE, Univ Montpellier, CNRS, EPHE, IRD, 34090 Montpellier, France; Department of Ecology and Evolution, Biophore Building, University of Lausanne, 1015 Lausanne, Switzerland; Department of Biology, McGill University, Montreal, Quebec H3A 1B1, Canada; Department of Ecology and Genetics, Evolutionary Biology Centre, Uppsala University, 75236 Uppsala, Sweden; Department of Biology, University of Kentucky, 101 T.H. Morgan Building, Lexington, KY 40506, USA; National Herbarium of NSW, Botanic Gardens of Sydney, Mount Annan, NSW 2567, Australia; Evolution and Ecology Research Centre, School of Biological, Earth and Environmental Sciences, University of New South Wales, Sydney, NSW 2052, Australia

**Keywords:** flowering plants, floral morphology, reproduction, functional ecology, pollination, trait evolution, mating systems, sexual systems

## Abstract

- Flowering plants have many modes of sexual reproduction, notably varying from selfing to outcrossing, and from bisexual flowers to individuals with separate sexes (dioecy). These reproductive modes are associated with floral and life-history traits that have evolved together, making it difficult to interpret correlations between traits.
- We analysed variation in 21 traits related to flowers, pollination, mating, sexual systems and life history from 361 species representative of flowering plant diversity.
- Outcrossing was mainly found among long-lived, large-stature plants, but hermaphroditic outcrossers and dioecious species appeared as largely non-overlapping strategies in the trait space. Level of floral investment was the main difference between these strategies, with dioecious species having generally smaller, less rewarding flowers, a pattern that also occurred in biotically pollinated species.
- This multi-trait study shows that pollination can be achieved in many, often contrasting, ways. Despite extensive variation in reproductive traits, dioecy stands out as being linked to floral traits primarily, while correlations with lifespan and dispersal traits appear spurious. We provide a conceptual framework based on lifespan, floral investment, and sexual separation that can be used to integrate pollination, reproduction and plant growth in future research on plant evolution and ecology.

## Introduction

Angiosperms (flowering plants) are by far the most species-rich group of plants today. Their reproductive organ, the flower, presents an exceptional diversity. It is thought this diversity has evolved to deal with the challenge of fertilization, for which flowering plants have to rely on external vectors, either animals or abiotic factors (wind, water), to mate with other individuals. Sex is considered to have evolved to facilitate the recombination of genetic material, allowing species to adapt to their environment (Maynard Smith, 1978), and recombination is more effective when individuals mate with other individuals (outcrossing) than with themselves (selfing) (Nordborg, 2000). Indeed, most plant species rely on outcrossing to produce offspring (Igic & Kohn, 2006), and it has been identified as a major driver of plant diversification (Goldberg et al., 2010; Vamosi et al., 2018; Hernández-Hernández & Wiens, 2020; Helmstetter et al., 2023; Anderson et al., 2023).

Most flowering plant species have bisexual flowers (here referred to using the botanical term ‘monocliny’), which is probably the ancestral state of the clade (Sauquet et al., 2017). In these species, outcrossing is often facilitated by genetic self-incompatibility (GSI), but many morphological features also exist that are thought to favor outcrossing, such as the differential maturity of pistils and stamens (dichogamy, heterodichogamy), and the disposition of pistils and stamens in the flower (herkogamy, distyly; Barrett, 2002). Dicliny, in which stamens and pistils occur in different flowers, could also have evolved to avoid selfing. The strongest separation of sexual functions is found in dioecious plants, which have separate ovule– and pollen-producing individuals (females and males), rendering selfing impossible. Dioecy has arisen frequently across the angiosperm tree of life but is found in only a small minority of species (Renner, 2014). It has been argued that dioecy is a “second-rate” outcrossing mechanism compared to GSI, as individual plants can only reproduce through one sexual function in dioecy, which would lead to lower fitness, all else being equal (Barrett, 2010, 2013). Another sexual system, monoecy, in which individuals bear unisexual flowers of both kinds, has been much less studied although it is about as frequent as dioecy (Renner, 2014). Finally, some sexual systems are characterized by a combination of unisexual and bisexual flowers, such as gynodioecy and andromonoecy, but they are much rarer and seem to be more restricted to particular areas or plant families (Bawa & Beach, 1981; Torices et al., 2011; Dufay et al., 2014; Renner, 2014).

The role that selfing avoidance has played in the evolution of dioecy has intrigued naturalists for more than a century (Darwin, 1884; Charlesworth & Charlesworth, 1978). Other floral traits could promote outcrossing in monoclinous species, possibly with fewer negative side-effects (Anderson et al., 2023). Comparing the traits of monoclinous and diclinous species, and their variation among mating systems, could shed light on the evolutionary mechanisms that led to the extraordinary diversity of flowers we find in nature. Furthermore, given the importance of mating and floral traits in evolution (Barrett, 2002; Vamosi et al., 2018), it is surprising that they have been largely neglected in the study of ecosystem functioning (E-Vojtkó et al., 2020). Traits and trait combinations evolve in the ecological context of the plants that possess them (de Jong & Klinkhamer, 2005), so by summarizing floral trait variation we also aim to facilitate the inclusion of floral traits in ecological studies.

Variation in mating and sexual systems is closely linked to differences in other floral traits. The most clear example probably is the “selfing syndrome”, in which species that mainly reproduce through self-pollination experience a reduction in flower size and attractiveness (showiness, scent, rewards) (Sicard & Lenhard, 2011), presumably because the selection pressure to maintain pollinator attraction has disappeared. In the case of dioecy, several flower traits have been hypothesized to favor its evolution, based on large-scale correlations (Renner & Ricklefs, 1995; Vamosi et al., 2003). Theory based on resource allocation predicts that small flowers resulting from less investment into pollinator attraction could favor the evolution of dioecy (Charnov et al., 1976). Indeed, non-attractive, wind-pollinated flowers have been thought to be associated with dioecy for this reason. However this hypothesis falls short in tropical rainforests where many dioecious species are found but pollination by wind is rare (Bawa & Opler, 1975). It has also been proposed that the type of biotic pollinator might influence the evolution of dioecy, but again the predictions are contrasting. In species that have many small bisexual flowers that are pollinated by generalists, the transfer of pollen between flowers of the same plant (geitonogamy) would lead to selfing, or to pollen discounting in self-incompatible species, in which case dioecy would be advantageous (Bawa, 1980). However there are issues with this idea – generalist pollinators might be too unreliable for dioecious species as they might visit one sex preferentially; and many dioecious species have specialized pollinators (Renner & Feil, 1993). An alternative hypothesis is that small flowers have less potential to diverge between the sexes due to sexual selection, reducing the possibility that pollinators prefer one sex (Vamosi & Otto, 2002). Strikingly, monoecy and its associations have been much less studied, although some hypotheses linking it to wind pollination have been put forward (Friedman & Barrett, 2008; Faegri & Van Der Pijl, 2013). Monoecy is often not considered in its own right, but instead as a form of hermaphroditism alongside monocliny (e.g. Maynard Smith, 1978; Charlesworth and Charlesworth, 1978, but see Bawa and Beach, 1981; Cronk, 2022).

Floral traits such as flower shape and colour are mostly considered to result from the adaptation to different pollinators (Faegri & Van Der Pijl, 2013; Rosas-Guerrero et al., 2014). These traits could also influence the rate of outcrossing because they affect interactions with pollinators (Wessinger, 2021). It has been noted, for example, that wind-pollinated species tend to either preferentially rely on outcrossing or on selfing, while biotically pollinated plants present a more or less continuous distribution of outcrossing rates including intermediate ones (Aide, 1986; Vogler & Kalisz, 2001). Bilateral symmetry (zygomorphy) is thought to enhance pollen transfer by constraining pollinator position or by attracting more specialized pollinators, and could thus lead to increased outcrossing compared to species with radially symmetric (actinomorphic) flowers (Citerne et al., 2010).

Non-floral traits have also been shown to be linked with mating and sexual systems. Variation in mating system (predominant outcrossing to predominant selfing) is associated with lifespan and plant size. Selfing is mainly found among smaller, annual species because they rely heavily on the reproductive assurance selfing provides, while large, long-lived species with multiple opportunities for reproduction are thought to suffer more from inbreeding depression and thus typically reproduce through outcrossing (Scofield & Schultz, 2006; Petit & Hampe, 2006). The same reasoning could apply to sexual systems, with dioecy as an obligate outcrossing system that is relatively common among trees (Renner & Ricklefs, 1995; Vamosi et al., 2003). In addition, more efficient pollen transfer through the avoidance of geitonogamy could favour dioecy over other outcrossing strategies in large plants (Holsinger, 1988; Harder & Wilson, 1998).

Last, dispersal traits could be related to mating and sexual systems. Species with long-distance dispersal would benefit from the ability to self-fertilize as mate availability is usually low in newly established areas (Baker, 1955; Rodger et al., 2018). However, more recent theory provides the contrasting prediction that under spatial heterogeneity in pollen limitation, outcrossing-dispersal vs selfing-nondispersal syndromes can evolve (Cheptou & Massol, 2009; Massol & Cheptou, 2011). For dioecious species, in which seeds are produced only by half of the individuals, more efficient dispersal by animals would decrease the local competition among seedlings in the vicinity of female plants (Heilbuth et al., 2001). An alternative explanation for the association of biotic dispersal and dioecy is that the uneven costs for reproduction would lead to sexual specialization (Bawa, 1980; Charnov, 1987).

Floral and other life-history traits are thus inextricably linked with mating and pollination, either directly or indirectly. As any given species possesses a multitude of traits, they act together to influence its evolutionary and ecological success (Anderson et al., 2023). However basing our understanding of trait interactions on correlations between pairs of traits could lead to misinterpretation. Dioecy, which, as any outcrossing mechanism, is predominantly found among trees, could be statistically correlated to other features of trees by coincidence, and without any direct link with floral and dispersal traits. For example, fleshy fruits have been correlated with dioecy (Vamosi et al., 2003), but this could simply be a by-product of the fact that trees in general can invest more resources in dispersal than small species for which dispersal is mostly unassisted (Thomson et al., 2018). Similarly, flower characteristics might be different between trees and herbs, as suggested by the observation that zygomorphy is characteristic of several large, mainly herbaceous families such as the Orchidaceae or Gesneriaceae (although this has not explicitly been tested to our knowledge). To disentangle covariation in sexual, mating, floral, and other life-history traits, they must be studied conjointly. Doing so will help uncover what traits are most closely associated with outcrossing itself, as well as with the different ways to promote outcrossing.

Some of the traits associated with variation in outcrossing rates, such as lifespan, plant size and dispersal, are classically considered in plant ecology to define “plant strategies” (Grime, 1974). These strategies determine local species composition based on the plant’s abilities to cope with competition, disturbance and stress. They are, as yet, not explicitly linked to pollination, but it is likely to play a role in plant community composition, as interactions with pollinators influence species coexistence through seed set and seed quality. While it is becoming clear that floral traits vary in ways not captured by vegetative ones (E-Vojtkó et al., 2022), they are currently not available at a large scale (Kattge et al., 2020), thus limiting their use in functional ecological studies. Describing the main axes of variation in floral traits will allow to identify traits that could help ecologists understand plant community assembly.

In this study we aim to describe trait variation at the angiosperm level. As much of the diversity in angiosperms occurs in regions in which the flora is only sparsely studied, we focused on morphological traits that are usually documented in botanical descriptions. We included mating systems (from selfing to outcrossing), for which we could gather information from the scientific literature, sexual systems (monocliny, monoecy and dioecy), as well as other traits which are reliably indicated in the botanical literature. This led us to a dataset of 21 traits in 361 species. We used these data to examine how sexual systems, flower morphology, and pollination modes collectively influence variation in plant mating. We built reproductive trait spaces to (1) determine the extent to which floral traits and pollination modes co-vary at the angiosperm level, (2) explore how mating and sexual systems are distributed across the main axes of variation and (3) establish whether major reproductive strategies can be characterized among flowering plants.

## Materials and Methods

### Data collection

We collated trait data for angiosperm species using the PROTEUS collaborative database (Sauquet, 2019). Our aim was to obtain a representative sample of the angiosperm diversity. We started with species from the angiosperm-wide dataset of López-Martínez et al. (2023) for which reproductive information (e.g., outcrossing rates, self-compatibility, dioecy) was available. Then, we expanded the species sampling by adding at least one species from each family with more than 100 species. We added more species for the most species-rich families (e.g., Asteraceae, Orchidaceae), choosing species that belong to the main clades of these large families to best represent their diversity. This process led to an initial set of 363 species.

We selected a list of traits based on prior knowledge of how reproduction-related traits influence evolutionary success (Helmstetter et al., 2023; Anderson et al., 2023). These traits were selected primarily to encompass the main aspects of angiosperm reproduction and included those related to mating system, sexual system, floral morphology (flower sex, ovary position, flower colour, flower size, flower symmetry), dispersal distance/mode and pollination mode. We also included several vegetative traits related to growth form and lifespan, which can also be related to reproduction (see Introduction). For each trait, detailed scoring instructions were followed; for traits already in PROTEUS, we used the instructions from Sauquet et al. (2017) and Schönenberger et al. (2020), whereas for newly added traits, we compiled instructions (Notes S1) based on previously available guidelines (Perez-Harguindeguy et al., 2013; Cardoso et al., 2018). Seed mass was added outside PROTEUS as the species mean according to the Seed Information Database (SER et al., 2023). Traits were scored at the species level (cf Sauquet & Magallón, 2018); intra-specific polymorphisms were scored as such, and only traits that could be reliably scored for a species were used.

To compare our results with those derived from vegetative and seed traits, we also analysed a data set of six plant traits (leaf area, leaf mass per area, leaf nitrogen per mass, diaspore mass, stem specific density and plant height) for *>* 45,000 species (Díaz et al., 2016; Díaz et al., 2022).

### Trait encoding

To facilitate downstream analyses, we modified the initial trait encoding to create a tractable and interpretable set of traits (Table S1). For qualitative traits, we reduced the number of states to between two and four for comparisons to be informative. In some cases we split an initially complex trait into multiple different ones for easier interpretation of results. For example, habit was recoded into three binary traits: woodiness, climbing and aquatic. Similarly, the original sexual system (including monocliny, dioecy, monoecy, gynodioecy, andromonoecy, etc.) was split into two binary traits: (1) flower sex coded as unisexual vs bisexual and (2) sexual system, coded as monomorphic (including monocliny, monoecy, andromonoecy and gynomonoecy) vs dimorphic (including dioecy, androdioecy and gynodioecy). For quantitative traits, if several values were available for a species/trait combination (either several measurements or indication of minimum and maximum values), we used their mean. The outcrossing rate was transformed into a qualitative trait “mating” using three bins: selfing (*<* 0.2), mixed mating (0.2 *−* 0.8) and outcrossing (*>* 0.8). Although approximate, this classification reflects the observation that mixed-mating species often have large temporal and/or spatial variations in realised outcrossing rates, whereas in predominantly selfing and predominantly outcrossing species outcrossing rate are often more stable (Whitehead et al., 2018). This allowed us to combine species with a quantitative estimate of outcrossing rate with those for which only a qualitative classification was available (phenotypic mating system, self-incompatibility and dioecy, see Table S1).

We encoded the qualitative traits in two ways to facilitate different downstream analyses. The first had one variable per trait with as many values as there are states in the trait, plus separate values for cases in which the trait was polymorphic for a species (e.g. the trait woodiness has the states: woody, herbaceous, woody herbaceous). We refer to this encoding as the “original” data set. In the second, qualitative traits were encoded using a one-hot approach, where each category of a trait is treated as a distinct binary variable (e.g. woodiness is split into two variables, each with two states: (1) woody vs. non-woody, (2) herbaceous vs. non-herbaceous). While one-hot encoding may introduce some redundancy (e.g., most species that are herbaceous are not woody and vice-versa), it is an alternative way of dealing with polymorphic states while keeping the relations between values. For example, in the original encoding, a species that can be both woody and herbaceous is assigned to a separate category with no explicit relation to other woody and herbaceous species, while in the one-hot encoding, such a species is similar to both herbaceous and woody species.

For visualization purposes, we chose to divide species in five categories. Among species with bisexual flowers (monocliny), species were assigned according to their predominant mating system (selfing, mixed mating, and outcrossing). The two other categories concern species with unisexual flowers, among which we distinguish dioecious and monoecious species. The few species with both bisexual and unisexual flowers were considered monoclinous for this purpose, as usually, unisexual flowers are not found in all populations (e.g. in gynodioecy) or only represent a small fraction of the flowers. Species having both monoecious and dioecious populations were labeled according to their major sexual system, if this information was available, or not labeled. Again, this labeling was only used to aid the interpretation of the figures and data, and not in the multivariate analyses.

### Filtering, transformations and missing data imputation

To limit the impact of missing data on our analyses, traits were removed if more than 50% of values were missing in the original data set. Likewise, species were removed if more than 50% of their traits were unknown. We log-transformed quantitative traits to conform better to normality expectations, except for fusion of ovaries that was coded as a proportion. We also centered and scaled these variables to limit potential biases caused by using traits with different units of measurement.

After filtering, many of the traits still contained missing data. We conducted imputation with the ‘missForest’ R package (Stekhoven & Bühlmann, 2012) to determine how this affected distances between species. We followed the approach outlined in Debastiani et al. (2021). Briefly, a pairwise phylogenetic distance matrix (see below for how the phylogenetic tree was generated) containing all species was decomposed to a set of eigenvectors using the function *PVRdecomp()* from the ‘PVR’ R package (Santos, 2018). The first 10 eigenvectors were added to the trait data set as additional, complete traits to conduct imputation of missing data with the *missForest()* function.

We then examined pairwise correlations between traits in our original data set using different approaches depending on the type of traits being compared. For qualitative vs qualitative trait comparisons we calculated Cramer’s *V* using the *cramerV()* function in the R package ‘rcompanion’ (Mangiafico, 2025). For other comparisons we performed ANOVA (qualitative vs quantitative) or Pearson’s correlation coefficient (quantitative vs quantitative). We then used hierarchical clustering based on Gower’s distance to group traits together with the *hclustvar()* function from the R package ‘ClustOfVar’ (Chavent et al., 2017).

### Trait spaces

To build trait spaces, we first calculated pairwise distance matrices among species using Gower’s distance (Gower, 1971) with the function *daisy()* from R package ‘cluster’ (Maechler et al., 2022). Gower’s distance was used because it can deal with missing data and mixed data types (e.g. qualitative and quantitative). We then performed principal coordinates analysis (PCoA), a dimensionality reduction approach used to summarize similarities in the data, on the resultant distance matrix. We used the *pcoa()* function of the R package ‘ape’ (Paradis & Schliep, 2019) to generate a set of orthogonal eigenvectors and their associated eigenvalues. For the one-hot data set, we used the *wcmdscale()* function in the R package ‘vegan’ for the PCoA, and fitted the individual traits on the resulting trait space with the function *envfit()*. We also built an additional trait space using the vegetative traits in the Díaz et al. (2022) data set. To do so we first removed those species that had information for fewer than four of the six traits. This ensured distances could be calculated between all species pairs while increasing computational feasibility and accuracy of distance calculations. To compare our data set with Díaz et al. (2022) we extracted the 159 species common to both datasets and built the two corresponding trait spaces using only these species.

We quantified the “quality” of dimensionality reduction of the resultant trait spaces using the method outlined in Mouillot et al. (2021). Briefly, the difference between the initial distance matrix and the distance matrix after dimensionality reduction using PCoA was examined. High-quality trait spaces are those in which a reduced number of PCoA dimensions accurately represents initial distances among species, thus indicating high redundancy among traits. Quality was quantified using the area under curve (AUC) metric relating the increase in quality with increasing number of retained PCoA axes. This approach also provides an indication for how many axes are sufficient to summarize the variation in the initial dataset.

PCoA is a linear dimensionality reduction approach that does not account for more complex non-linear patterns. Therefore we also used an alternative dimensionality reduction approach, Uniform Manifold Approximation and Projection (UMAP, McInnes et al., 2020), to visualise non-linear patterns in our data. UMAP is based on manifold learning techniques and can be used to assess patterns at local and global scales simultaneously, depending on the size of the neighbourhood (‘n neighbours’) chosen. We applied UMAP to our Gower’s distance matrix calculated using the original data set, using the *umap* R package with the default configuration. We set the number of components (dimensions) targeted to two and varied ‘n neighbours’ (10, 25, 50, 100) to test the effect of changing this parameter on the distribution of species in the space. A small number of neighbors creates tight clusters based on local similarities, while a large number emphasizes the global structure.

### Clustering

To help define reproductive strategies we assigned species to different groups using the partitioning around medoids (PAM) (Kaufman & Rousseeuw, 1990) clustering approach, as implemented in the ‘cluster’ R package (Maechler et al., 2022). This method takes a distance matrix as an input and is based on determining a set of medoids (points) that represent the structure of the data. Each species is assigned to a medoid, with the goal of minimizing the sum of the dissimilarities of each species to their closest medoid. PAM clustering was done using Gower’s distance matrices for both original and one-hot encoded data sets.

The number of clusters (i.e. value of *k*) was initially selected using silhouette width. This metric ranges from –1 to +1, where high values indicate that a point is similar to its cluster and different from neighbouring clusters. However it can be difficult to objectively determine the appropriate number of clusters that should be used to summarise the data set. To tackle this subjectivity issue, we examined how cluster membership changed as values of *k* were changed using Sankey plots, a type of flow diagram. We then identified groups of species that consistently grouped together as *k* was increased from *k* = 2 to *k* = 7. We took the largest groups until the total number of species reached 80% of the species in our data and considered these as ‘robust groups’.

### Phylogenetic tree and simulated data sets

We built a phylogenetic tree among our species using V.PhyloMaker2 (Jin & Qian, 2022). We used the default ‘GBOTB.extended.TPL’ backbone tree that was derived from a large phylogenetic tree of all seed plants (Smith & Brown, 2018) and built the tree using the default approach described as ‘scenario 3’ (Jin & Qian, 2022). Prior to building the tree we standardized genus and species epithets using the R package ‘TNRS’ (Boyle et al., 2013) and retrieved higher level taxonomy using ‘TNRS’ and another R package, ‘taxize’ (Chamberlain & Szöcs, 2013). 308 species were found in the backbone tree and the remaining 53 were successfully bound to the backbone tree, providing a complete phylogenetic tree for our data set.

To determine how phylogeny influences trait space for our set of species and traits we simulated trait data using the phylogenetic tree. To do so we first fitted trait evolution models to each trait in the original data set with missing data imputed. For quantitative traits we fitted Ornstein-Uhlenbeck (OU) models using the *fitContinuous()* function in the R package ‘geiger’ (Pennell et al., 2014) to estimate OU model parameters and root state values. For qualitative traits we fitted fixed-rate, continuous-time Markov (Mk) models using the *asr mk model()* function in ‘castor’ (Louca & Doebeli, 2018) to generate transition rate matrices and ancestral likelihoods for the root state. We allowed all transition rates to be different by using all-rates-different (ARD) models. We then used the estimated parameters and the phylogenetic tree to simulate new datasets with *rTraitCont* from the ‘ape’ R package (Paradis et al., 2004) and *sim.history* from ‘phytools’ (Revell, 2012). Traits were simulated independently and then combined into a single simulated dataset, from which we calculated distance matrices and ran PCoAs, as above. This process was repeated 1000 times.

## Results

After recoding and filtering, the final data set consisted of 21 traits (Table S1) for 361 species and 13% missing data (Fig. S1). With representatives from 260 of 416 families, and 61 of 64 orders (Fig. 1, Tables S2,S3; APGIV, 2016), our dataset included a broad range of angiosperm diversity. Imputing missing data only slightly changed Gower’s distances between species (Fig. S2, Mantel statistic r = 0.91), and so would likely have little impact on the following results. Using one-hot encoding also had a minor effect on distances, which remained highly correlated (Mantel statistic r = 0.95). In the following we used the “original” and the “one-hot encoded” data; the imputed data were only used for comparisons among data sets and in the simulations to test for phylogenetic effects.

**Figure 1:**
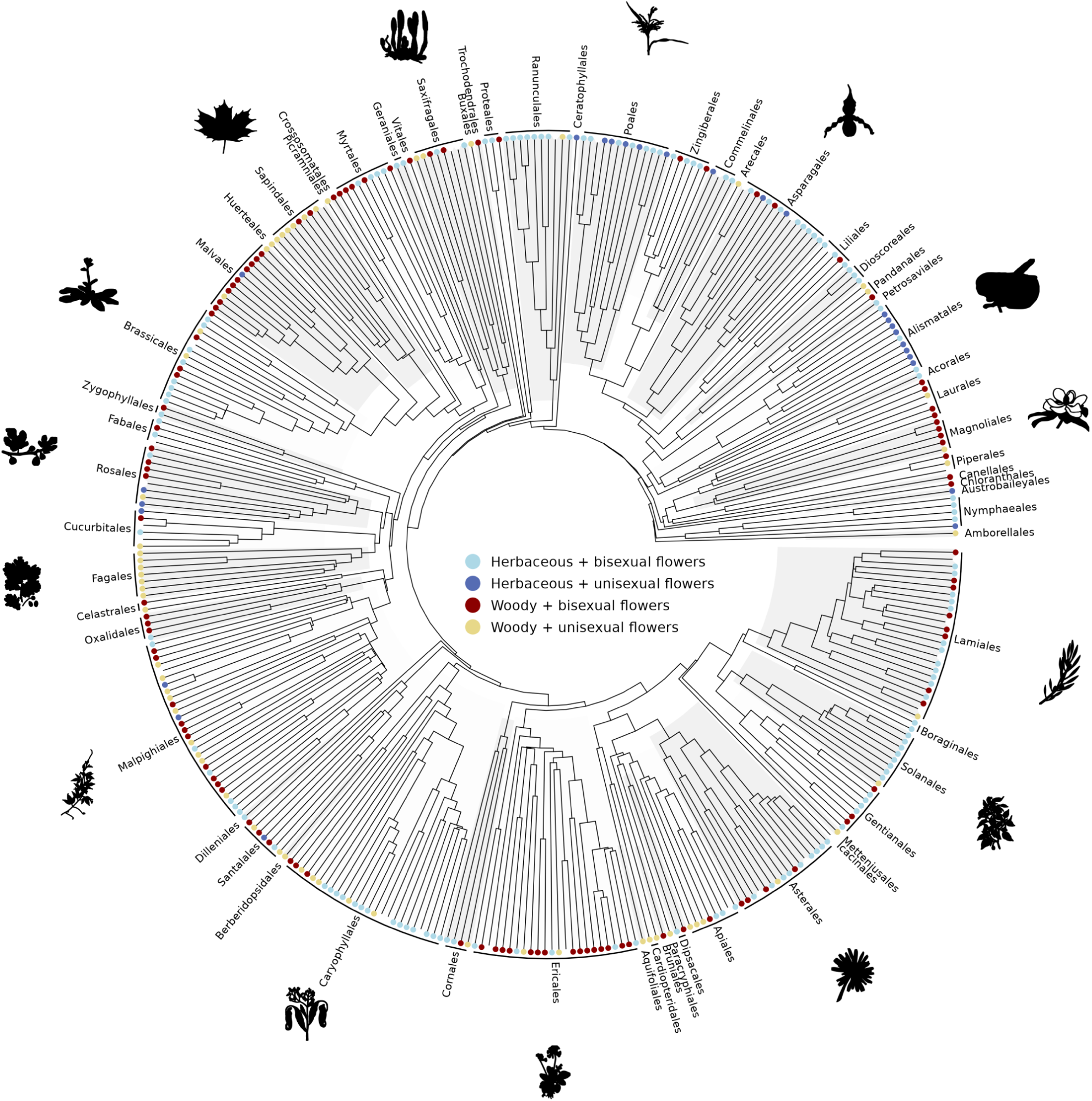
Phylogenetic tree representing the 361 angiosperm species used in our analyses. Tips are labelled with coloured circles indicating the combination of woodiness and flower sex each species possesses. Species without data or those that are polymorphic for either trait are left blank. Orders that are represented in our set of species are highlighted around the outside of the tree alongside a selection of species’ silhouettes from phylopic.org.

### Reproductive trait space: quality and dimensions

The first two PCoA axes of our trait space explained 30 to 34% of the variation depending on the data encoding used (one-hot encoding: Fig. 2a; original dataset: Figs. 3a, 4a). To characterize this trait space, we first compared it to the one built with data for six vegetative and seed traits classically used in plant functional ecology (Díaz et al., 2022) from 7968 species. We examined the location of our species in this mainly vegetative trait space and found that they were scattered relatively evenly throughout (Fig. S3a). Then, using a set of 159 species shared between the two datasets, we found that the species’ distances based on the six traits from Díaz et al. (2022) were only weakly correlated with those based on the 21 traits in our study (Mantel statistic r = 0.285; Fig. S4).

**Figure 2:**
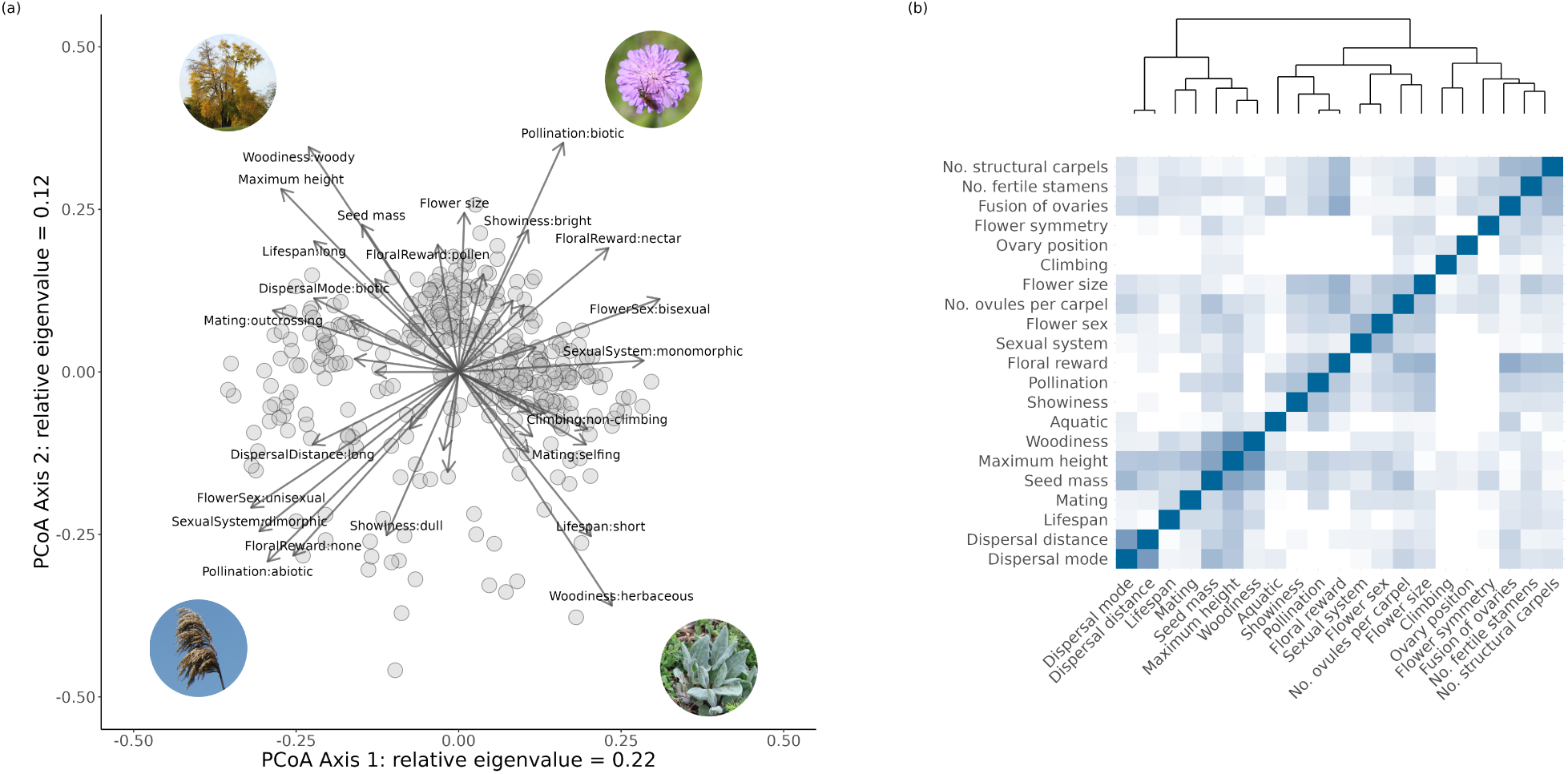
Multivariate analysis of flowering plant traits. (a) The trait space obtained with a principal coordinates analysis (PCoA) performed on the one-hot encoded data set. The (linear) vectors of each trait in the first two dimensions are indicated as arrows and the species are indicated as circles. Images representing aspects of the trait space are shown in the four corners of the plot; photographs by A. J. Helmstetter. (b) A heat map showing the strength of the correlations between pairs of traits, where darker blues indicate higher correlation. The correlation coefficients were calculated using the original encoding of the data, so correlation coefficients are presented as absolute values in all cases as the direction of correlation is not meaningful for categorical data (see methods). The dendrogram is derived from hierarchical clustering of all traits.

**Figure 3:**
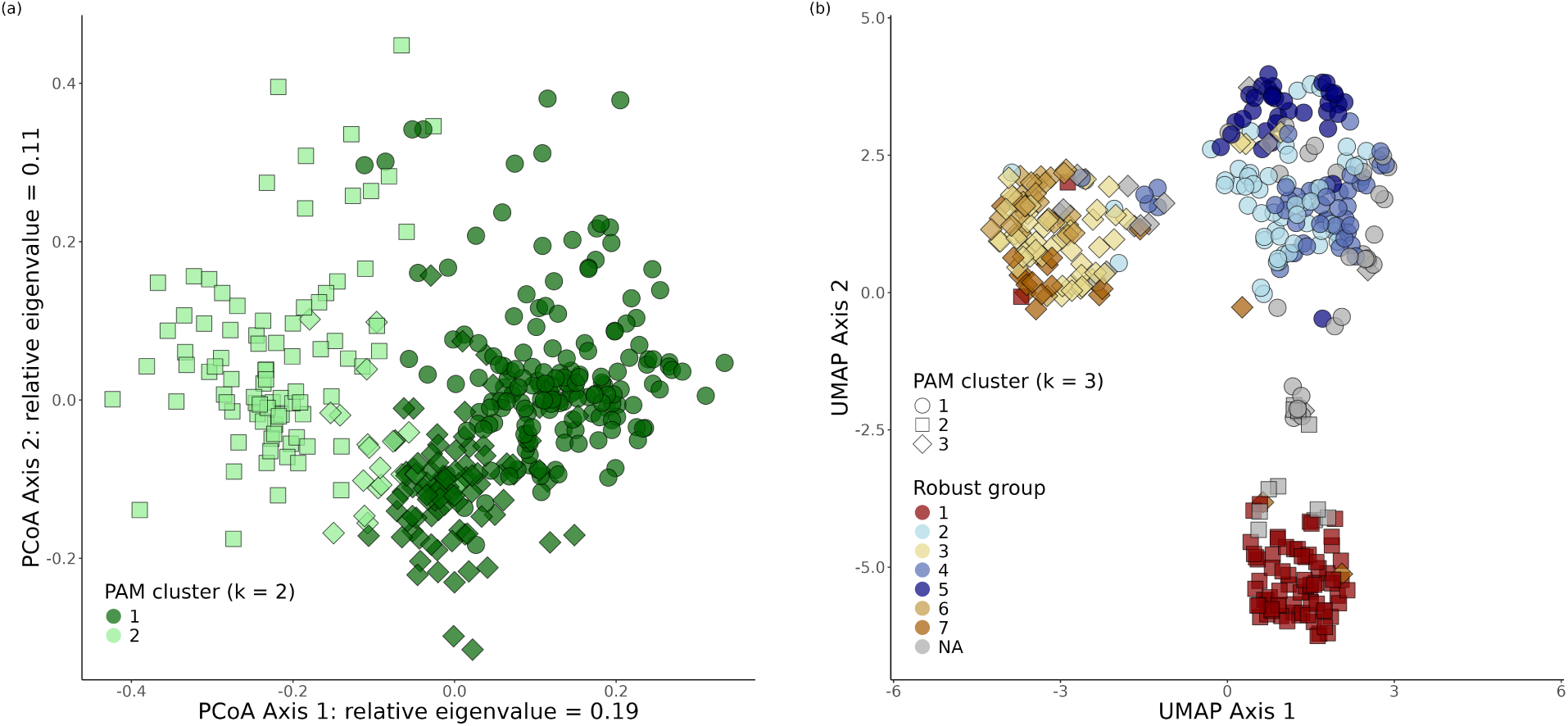
Trait spaces and clustering of angiosperm species. (a) the position of species on the first two axes of a principal coordinates analysis (PCoA) on the original data set. Points are coloured by cluster membership derived from clustering using the partitioning around medoids (PAM) method when *k* = 2 (number of clusters). (b) the distribution of species along the first two axes of a UMAP (uniform manifold approximation) analysis with a neighbourhood size of 10, showing final-scale structure in the data. Points are coloured by robust group (see Figure S12 for further details). Point shape in both panels indicates their PAM cluster assignment when *k* = 3. Circles are mostly herbaceous species with bisexual flowers, squares are species with unisexual flowers and diamonds are woody species with bisexual flowers.

**Figure 4:**
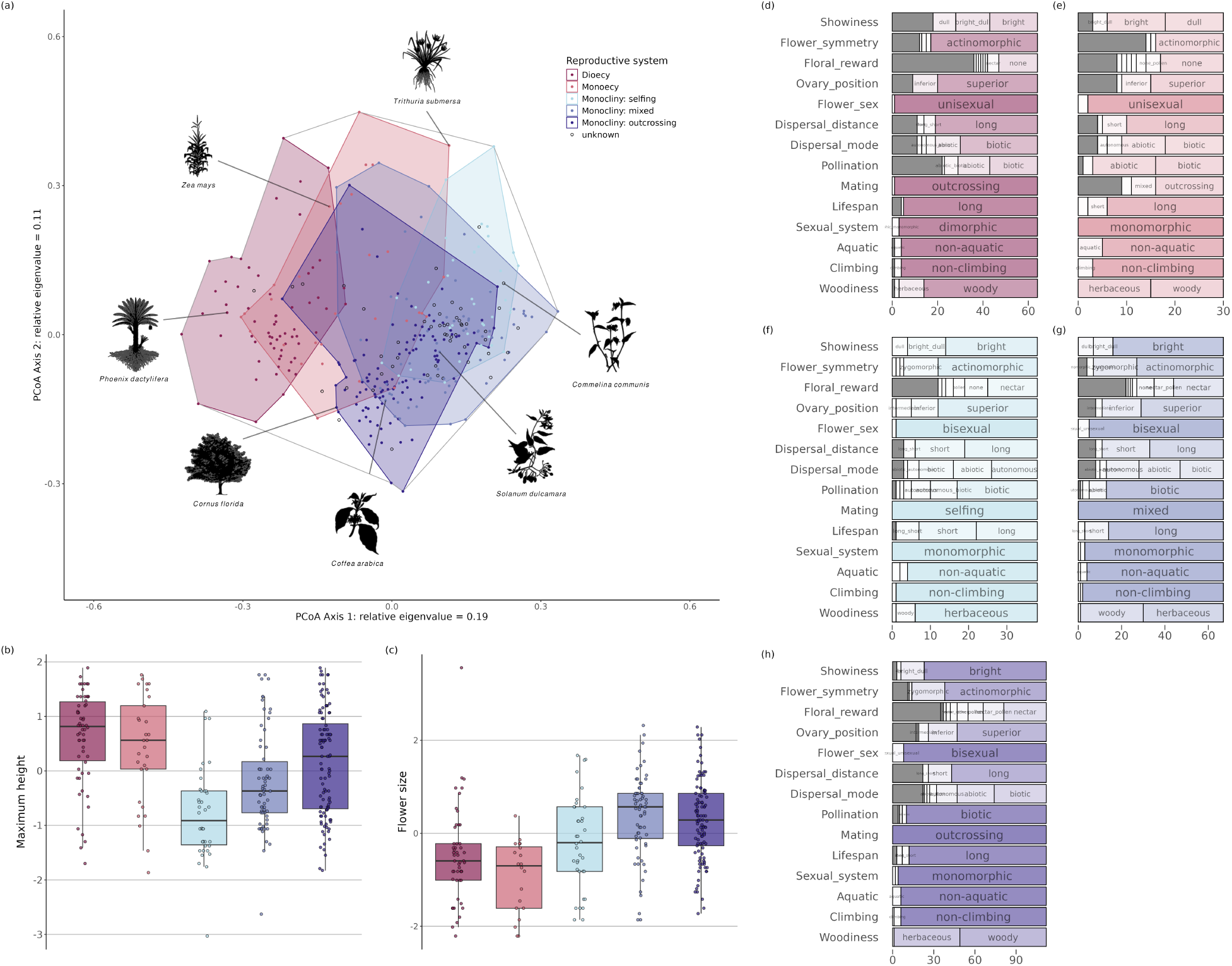
Sexual and mating systems in the angiosperm reproductive trait space. Panel (a) shows the position of 361 angiosperm species on the first two axes of a principal coordinates analysis (PCoA) on the original data set. Colours indicate the reproductive mode reduced to five categories for visualization. Boxplots show the distribution (after log transformation and scaling) of values for two traits per cluster: (b) maximum vertical height and (c) flower size. Points represent the values of species within each cluster. The stacked barplots (panels (d-h)) show the frequency of states for 14 categorical traits for each of the five reproductive modes: (d) dioecy, (e) monoecy, (f) monoclinous selfers, (g) monoclinous mixed mating species and (h) monoclinous outcrossers. Sections representing states with high frequencies are labelled and the dark grey sections correspond to missing data. Species silhouettes in panel (a), used to illustrate the diversity of species in our data set, are taken from phylopic.org. All are public domain except for *Cornus florida*, which is attributed to Gabriela Palomo-Munoz (CC BY 4.0 license, https://creativecommons.org/licenses/by/4.0/ – no changes were made).

We then used he approach of Mouillot et al. (2021) to calculate statistics to allow us to compare our trait space to others. The AUC criterion, which indicates how well the first *n* axes summarize the total variation in the data set, indicated that to get a good representation of our trait space, we must keep more dimensions than for the data set derived from Díaz et al. (2022) (Fig. S3b). This was equally the case when comparing our results to those of the other trait spaces analyzed by Mouillot et al. (2021). This means that, in our data set, traits were weakly redundant and most of them contribute small but significant amounts of variation that cannot be reduced to variation in other, more structurally important traits. We did not see notable differences in the trait space quality analyses between our different data sets (original, imputed, one-hot; Fig. S5).

In order to test for phylogenetic inertia, we performed ancestral state reconstruction with the original dataset to estimate parameters that were used to simulate new data sets where all traits evolved independently. As expected, distance matrices from these simulated data sets were not correlated to the real data sets (Mantel statistic r = 0.03 using an example simulated data set) and maximum distances were greater when using real data (Fig. S2). When we ran PCoA on simulated data sets we found that the first three to four axes explained substantially less variation than in the real data sets (Fig. S6). Thus, at least the first three, perhaps four, PCoA axes from the analysis of our original data set are due to co-evolution of traits rather than neutral phylogenetic co-occurrence.

### Trait covariance

The first four axes of the trait space were correlated with several general life-history and floral traits (Table S4). Among the most important traits we found woodiness, lifespan, seed mass and plant height, and they were co-distributed along a diagonal in the 2D trait space defined by the first two PCoA axes (Fig. 2). This same diagonal also corresponded to mating system variation, from selfing, which was associated with small size and short life span, to outcrossing, which was associated with woodiness, large size and long life span. Many flower and pollination-related traits contributed to variation that was orthogonal to the size/lifespan diagonal, including flower sex, floral reward, biotic vs. abiotic pollination, showiness and plant sexual system. The third PCoA axis, representing 9% of the variation, was mainly associated with dispersal (mode and distance), and the fourth axis, representing 7% of the variation, with ovary position and flower symmetry (Fig. S7).

We examined correlations among traits (Fig. 2b) and found two main groups, reflecting patterns in the trait space. One group contained vegetative, mating and dispersal traits, and the other contained flower morphology, pollination and sexual system. Among the first group of traits, lifespan, height, seed mass, woodiness and mating were most strongly clustered, while dispersal traits fell into the same group but were more weakly associated. Among the floral traits, two subgroups could be identified, one mainly related to pollination and attraction (flower size, showiness, reward), and another subgroup concerning the reproductive organs themselves (carpel and stamen number, position and fusion of ovaries, flower symmetry). Flower sex and sexual system, although clearly concerning the reproductive organs, clustered with the pollination and attraction traits, not the other carpel and stamen traits, which instead clustered with flower symmetry.

### Major reproductive strategies

Partitioning Around Medoids (PAM) clustering (Figs. 3 & S8) of the species based on their trait-based distances pointed to the existence of two major groups (light green and dark green points in Fig. 3a). These were well separated in the first two dimensions of the trait space, and can be predominantly characterized by species with bisexual flowers vs species with unisexual flowers. When the number of clusters was increased to three, the cluster of unisexual species remained (Squares in Fig. 3) while the cluster of species bisexual flowers was split into a herbaceous (circles) and a woody (diamonds) group (Figs. 3 & S9). Such a structure in three major groups was also revealed using UMAP (Fig. 3b; Fig. S10), which is based on an alternative decomposition approach that allows the visualisation of non-linear and local patterns.

We sequentially increased the number of PAM clusters up to seven (Fig. S8) and tracked whether groups of species stayed together in the clusters (“robust groups”) or not (Fig. S11). The species with unisexual flowers remained a markedly stable group (Fig. S12d), especially the woody dioecious species; the unisexual species that did not remain in this group were mostly monoecious and herbaceous. The species in the bisexual clusters were further split into three robust groups each: among the woody species, one group with smaller, rather dull flowers stands out (Fig. S12c,j), while among the herbaceous species, a distinct robust group with zygomorphic flowers appears (Fig. S12h). Thus, traits such as zygomorphy, flower size and dispersal mode, are characters that play different roles depending on the context of other traits (mainly woodiness). Remarkably, mating system was not clearly associated with one of the clusters, even among herbaceous species where most variation in this trait is found. A total of 55 species did not group with other species throughout the clustering process and thus were not assigned to robust groups. Many of these were found in distinct, sparsely populated areas of the trait spaces – in the top centre of the PCoA plot (Fig. 3a) and in the middle of the UMAP plot (Fig. 3b). In general, these were long-lived, herbaceous species with small, dull, abiotically-pollinated flowers, some of which were aquatic.

### Mating and sexual systems

In all analyses, variation in reproductive modes was encoded by three separate traits (mating, flower sex, sexual system; Table S1). To facilitate interpretation we plotted the original sexual systems (dioecy, monoecy, monocliny) and the mating systems of monoclinous spaces back on the trait space. This allowed us to visually assess that dioecious species occupied an area in the trait space that was largely distinct from the area occupied by the monoclinous species (Fig. 4a). Monoecious species were found between these two sets and overlapped substantially with both monoclinous and dioecious species, occupying a large area. Among the monoclinous species, the variation in mating system was associated with a gradual shift in the associated traits. We found that predominantly outcrossing and selfing species shared a large overlapping area in the trait space, despite being at opposite ends of a major axis of variation.

There was a gradual increase in average size from selfing to outcrossing monoclinous species, whereas dioecy and monoecy seemed more restricted to large-stature plants (Fig. 4b). Similarly, while monoclinous selfing species had on average smaller flowers than outcrossers, average flower size was even smaller among dioecious and monoecious species (Fig. 4c; see Fig. S13 for all quantitative traits). The associated categorical traits (Fig. 4d-h) confirmed the trait associations visible in Fig. 2. Unisexual flowers were almost always actinomorphic, while there was no notable difference in flower symmetry among the monoclinous species with different mating systems. Abiotic pollination was found more frequently among species with unisexual flowers, as was the absence of a floral reward. However, purely abiotic pollination was not the major mode of pollination among dioecious species, and its frequency was higher (about 45%) among monoecious species.

## Discussion

We here present an angiosperm-scale synthetic analysis of the plant traits associated with mating and sexual system variation. We compiled information on 21 traits, combining classical life-history traits (plant size, life form, dispersal), with those relating to flowers, pollination and reproductive modes. Our study is based on 361 species that were chosen to represent the angiosperm diversity, including species from more than 50% of the families and nearly all orders (Fig. 1, Tables S2,S3).

Sexual and mating systems had markedly different distributions in the trait space (Fig. 4). Mating system variation was mainly correlated with variation in lifespan and size, as has been documented previously (e.g. Petit & Hampe, 2006; Salguero-Gómez et al., 2016). Sexual system variation, on the contrary, was linked to variation in floral and pollination traits. This pattern seems to be mainly driven by the contrast between dioecious and monoclinous species: among the species that are mainly outcrossing the dioecious species are those that have smaller, less rewarding flowers. The patterns we uncovered seem to be robust to trait encoding and multivariate method (PCoA, UMAP, clustering; Fig. 3), and the trait associations are much stronger than what would have been expected based on phylogeny alone (Fig. S6).

Along the lifespan-size axis, mating systems largely overlapped. Thus, outcrossing and selfing species could not easily be distinguished, and floral traits were only weakly discriminative at this scale. Selfing species tend to have smaller flowers than mixed-mating or outcrossing species, consistent with the observation that the ‘selfing syndrome’ often involves a reduction in corolla size (Sicard & Lenhard, 2011). However, the fraction of species with zygomorphic flowers, often interpreted as being the sign of high-precision pollination favoring outcrossing, was similar among selfing, outcrossing and mixed-mating species (Fig 4). Indeed, a transition to predominant selfing can arise in very different pollination contexts, e.g., wind-pollinated grasses (Burgarella et al., 2023), small-flowered herbs with generalist pollinators (Sicard et al., 2011), or in groups where specialist pollination syndromes have evolved (Rose & Sytsma, 2021). Furthermore, selfing is associated with higher extinction rates (cf Goldberg et al., 2010), although this might depend on associated traits (Zenil-Ferguson et al., 2019; Helmstetter et al., 2023; Anderson et al., 2023). Increased extinction in selfing lineages would limit the scope for co-evolution of multiple traits, which could explain why floral traits associated with selfing are specific to each clade.

Traits associated with sexual systems have been described at the level of angiosperms (Renner & Ricklefs, 1995; Vamosi et al., 2003) or regional floras, including species from many families (Bawa, 1980). Given the large number of traits thought to be correlated with sexual system variation (see Introduction), as well as the potentially contrasting effects of these traits (Anderson et al., 2023), it was rather surprising that dioecy and monocliny were very well separated in the trait space (Fig. 4). Floral and pollination-related traits accounted for most of the differences between dioecious and monoclinous species. By contrast, the correlation between dioecy and biotic dispersal seemed to be purely a by-product of the fact that dioecious species were more often woody, and woody species rely more often on biotic dispersal (Fig. 2).

Most studies on sexual systems contrasted dioecy with hermaphroditism, often either excluding monoecy or considering it as a particular case of hermaphroditism as, indeed, a monoecious individual can self-pollinate in the absence of an incompatibility mechanism (Bertin, 1993). Here we found that the traits of monoecious species were intermediate between those of monoclinous and dioecious species (Fig. 4). This is consistent with the idea that monoecy presents a lesser degree of sexual specialization than dioecy, and might serve as an evolutionary intermediate between dioecy and monocliny (Renner & Ricklefs, 1995). However, there was extensive variation in the traits associated with monoecious species (even though we sampled more dioecious than monoecious species), which could be related to the variation in the spatial organisation of unisexual flowers. For example, some species have inflorescences with both female and male flowers (e.g. *Hevea brasiliensis*, *Arum maculatum*) forming functionally bisexual floral units, while in others the female and male flowers are clearly separated (e.g. *Zea mays*). Monoecy has not been as intensively studied as dioecy (cf Cronk, 2022) but clearly warrants further consideration in its own right. Investigations into the drivers behind its evolution (e.g. resource allocation, adaptation to environmental conditions, sexual selection and interference (Willson, 1979; Bawa & Beach, 1981; Golenberg & West, 2013; Chen & Pannell, 2023)) and the spatial (and even temporal) organisation of flowers are ripe avenues for future research.

Much trait diversity remained unaccounted for by these large-scale associations, as was reflected by the fact that the first two axes of the trait space had relatively low eigenvalues compared to other datasets (Mouillot et al., 2021). This may have partly been due to those traits that vary mostly in subsets of species, but not consistently among the whole set of species (e.g. zygomorphy, dispersal mode; Figs. 3, 4, S12). Unraveling the trait associations in these subsets would require more data and will likely open new research questions. For example, why are zygomorphic flowers mainly found among herbaceous species, and why do they occur similarly among selfing and outcrossing species? Is there a correlation between floral traits and dispersal syndromes among tree species? The approach used in this study can be used to discover finer-scale patterns among species with shared sexual and mating systems, as well as at the level of plant orders or families.

### The intricate correlations of sexual systems

The evolution of sexual systems is thought to have been driven by selection for outcrossing, for resource allocation, for more efficient pollen transfer, or a combination of those (e.g Charnov et al., 1976; Charlesworth & Charlesworth, 1978; Bawa, 1980; Thomson & Brunet, 1990; Barrett, 2002; Käfer et al., 2017; Lesaffre et al., 2024). Our multi-trait approach can shed some light on the many hypotheses that have been invoked. First, it is obvious that dioecy can only occur in species for which outcrossing is favorable, and reproductive assurance of secondary importance only. Thus, both monoclinous outcrossing species and dioecious species are more common among long-lived, woody species in particular. Second, flower unisexuality and sexual dimorphism are above all correlated with other flower traits, and only weakly with dispersal (Fig. 2). It has been hypothesized that biotic dispersal would increase female fitness and thus favor the emergence of dioecy (cf Thomson & Brunet, 1990) or that biotic dispersal would be a way to counterbalance the fact that seeds are dispersed by only half of the individuals (Heilbuth et al., 2001; Vamosi et al., 2007). However, our results show that this correlation is likely due to the higher proportion of both dioecy and biotic dispersal among trees (Renner & Ricklefs, 1995; Thomson et al., 2018), although other confounding factors might play a role as well (Thomson & Brunet, 1990).

Among the flower traits that distinguish diclinous (i.e. both monoecious and dioecious) from monoclinous species are flower size, showiness, pollination mode and pollinator reward. In accordance with earlier studies, wind pollination was overrepresented among dioecious and monoecious species, and this mode of pollination is typically associated with smaller, dull flowers and the absence of a reward. Nevertheless, almost half of the dioecious and monoecious species we considered have been reported to be pollinated by animals. It should furthermore be taken into account that pollination modes and floral rewards are often inferred from floral traits and observed flower visitors instead of pollination assays, and thus might not be correct. For example, although palms (a family with a high incidence of dioecy; Nadot et al., 2016) had long been thought to be mainly wind-pollinated based on their often rudimentary flowers, current knowledge indicates that a large majority are pollinated by insects (Barfod et al., 2011; Henderson, 2024). So, the correlation of dioecy with small, unrewarding and rather inconspicuous flowers does not seem to be a side-effect of wind pollination, as it affects biotically-pollinated species alike. Rather, our results indicate that dioecy is more likely to evolve in small-flowered species, possibly because there are fewer resources to be shared between the female and male functions of those flowers (cf Charnov et al., 1976). It is possible that small, dull, insect-pollinated flowers have less potential to become sexually dimorphic (which could decrease pollination, Vamosi and Otto, 2002) than showy flowers, but given the often extreme differences between female and male flowers in small-flowered wind-pollinated species, this seems an unlikely explanation.

Of course, there were many exceptions to the general patterns. Among the most striking ones in our data set was a species of *Rafflesia*, the genus with the largest known flowers, which is dioecious. This species obviously also has some other traits making it difficult to compare to other plants – classifying it as herbaceous or woody would not reflect its unique, almost completely endoparasitic habit. Interestingly, parasitism is also overrepresented among dioecious species, but the reasons for this association are unknown (Renner & Ricklefs, 1995). We could not study this correlation here as parasitism is rare among plants, and our dataset included too few parasitic plants. Similarly, although a climbing growth form is more frequent and our dataset included several climbing species, there were nevertheless too few species to reliably detect any correlation between a climbing growth form and sexual system (cf Renner & Ricklefs, 1995).

### A conceptual framework for reproductive trait variation

Our results highlight the importance of including multiple aspects of plant reproduction when studying trait associations, as this allows a better apprehension of the most influential correlations. Of course, other traits contribute to the reproductive strategies of plants (Barrett, 2002; Barrett, 2003). For example, dichogamy, a difference in the timing of fertility of the pistils and stamens, could also lead to more effective pollen transfer between individuals. It has several variants (Bertin & Newman, 1993), including heterodichogamy, in which some individuals of a population are protogynous and others protandrous (Renner, 2001). Bertin and Newman (1993) rejected the hypothesis that dichogamy primarily evolved to avoid self-fertilization, and found intriguing differences in the traits associated with the several types of dichogamy. Similar patterns might exist for herkogamy and distyly. We suggest that these morphologies reflect a certain degree of separation of the sexual functions, similar to the sexual systems.

To allow the integration of additional traits into the reproductive strategies we have identified, we propose a conceptual reproductive trait space with three dimensions: lifespan, floral investment and sexual separation (Fig. 5). While lifespan is not strictly a reproductive trait we use it here because it is an easily measurable trait that summarizes how much a species relies on outcrossing: short-lived species can be either selfing or outcrossing, while long-lived species are almost always outcrossing. Floral investment is the allocation of resources into the production of a flower, with small, non-attractive and non-rewarding flowers on one end and large, attractive, nectar-producing flowers on the other. Sexual separation encompasses the sexual systems, from monocliny through monoecy to dioecy. The main strategies we characterized in this study form a 2D triangle in this 3D space, with (1) monoclinous, small-flowered annuals, (2) monoclinous, large-flowered trees and (3) dioecious small-flowered trees at the vertices. Most species will fall close to the surface of this triangle, which thus describes the most common strategies. Yet, this framework also incorporates less frequent strategies that occur at greater distances from this plane, such as wind pollinated herbs or showy annuals. Note that this does not imply that such strategies are either ecologically or evolutionarily less successful.

**Figure 5:**
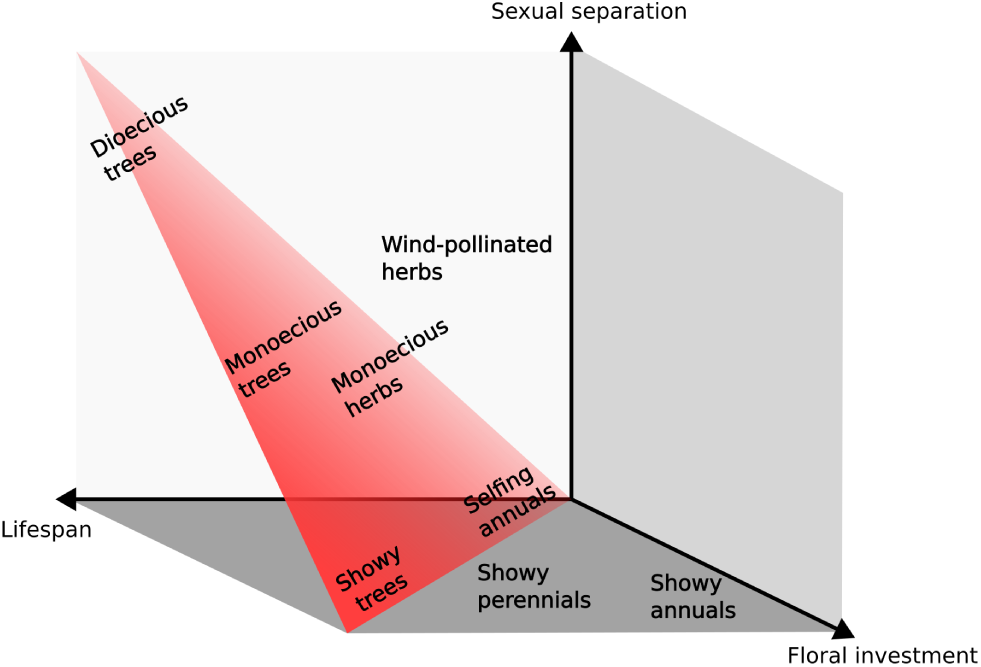
Conceptual representation of angiosperm reproductive strategies, defined by a 3D space with the most common strategies depicted on a 2D triangle. One main axis is lifespan, mainly related to outcrossing in the reproductive trait space: the longer the lifespan, the more a species will be outcrossing. The sexual separation axis includes the sexual systems included in our analysis (from monocliny to dioecy) but potentially also other ways of separating sexual functions, like dichogamy. The floral investment axis includes flower size and floral reward, and possibly flower longevity. Strategies that are not explicitly labeled as dioecious or monoecious are monoclinous.

In defining dimensions this way, we allow the framework to be expanded to traits we have not studied. For example, dichogamy and herkogamy could be situated somewhere on the sexual separation axis; would their associated traits differ markedly from, say, those of monoecious species? Where do systems that allow plastic allocation to female and male flowers fall (Lloyd & Bawa, 1984)? Other traits could include floral longevity (Stephens et al., 2024) and 3D structure (Van der Niet et al., 2010), as further measures of floral investment.

### Integrating floral and mating traits in the study of functional diversity

Just as plants have ecological strategies resulting from tradeoffs involved in competition, stress-tolerance and dispersal (Grime, 1974; Westoby, 1998), they have reproductive strategies. This term has been previously used to encompass life-history strategies that include diverse aspects of seed production and its contribution to population growth and persistence (Salguero-Gómez et al., 2016; Salguero-Gómez, 2017). We argue that these strategies should include floral and pollination traits: we have shown they account for substantial additional variation, and this variation could influence reproductive success and thus the composition of plant communities. On short timescales, pollination directly influences an individual’s fitness through the number of seeds produced (e.g., depending on pollen limitation; Ashman et al., 2004). Also, pollination can influence the fitness of offspring (if affected by inbreeding depression; Crnokrak and Roff, 1999), and could thus affect plants’ abilities to compete and cope with stress (Craig & Mertz, 1994; Cheptou et al., 2000; Petrone Mendoza et al., 2018). On longer timescales, pollination and reproduction affect genetic diversity and adaptive capacity, and thus most likely play an important role in long-term population and species survival (Burgarella & Glémin, 2017; Anderson et al., 2023). Hence, pollination could influence species’ ecological success, or maybe even its ecological strategy, although this remains to be investigated.

It is increasingly recognized that floral traits should be considered when characterizing functional diversity in angiosperms (E-Vojtkó et al., 2020). Here we aimed to represent angiosperm diversity by working with an original dataset chosen to represent all major angiosperm clades. This is complementary to the approach taken in other studies dealing with specific floras or datasets. E-Vojtkó et al. (2022) studied two datasets of European species (central Europe and Alps), and found that floral traits vary largely independently of vegetative traits. Despite their exclusion of trees and grasses, and their focus on a different set of traits (notably not including sexual system), the trait spaces they obtain are rather similar to our own. Lanuza et al. (2023) analyzed plant reproductive strategies in light of interactions with pollinators. Wind-pollinated species were therefore excluded, and tree species were generally underrepresented. Despite this, they also found that variation in selfing and outcrossing was only weakly correlated with variation in flower traits. The relative importance of floral traits in their trait space was higher than in our study, which could be due to the larger number of floral traits they studied (including style length, flower number per plant, quantity of nectar and pollen).

These studies, as well as ours, thus suggest that variation in pollination-related traits is largely independent of variation in vegetative traits, that are the main determinants of ecological strategies (Westoby, 1998). It should however be noted that the first axes of the trait spaces in these studies represent less than 50% of the total variation, leaving room for important correlations in other dimensions. At a first glance, as resources are limited, it might seem that investment in flowers would diminish investment in vegetative growth, and thus competition and stress-tolerance, and accordingly one would expect vegetative and floral investment to be negatively correlated. This intuition is however based on the assumption that growth and reproduction have contrasting roles in plants’ life cycles, by contributing to either growth or dispersal. This assumption might be false if, on the contrary, pollination and outcrossing influence competition and stress-tolerance (as discussed above). Thus, more studies are needed to compare, for instance, plant vegetative form and function (Díaz et al., 2016), fast-slow strategies (Salguero-Gómez et al., 2016), the leaf economics spectrum, the flower economics spectrum (Roddy et al., 2021), and other floral traits (Michelot-Antalik et al., 2025). In particular, it will be insightful to assess how floral and pollination traits vary in the context of plant communities, and how vegetative traits such as the specific leaf area, useful to distinguish stress-tolerant and competitive plants, interact with floral traits to influence species composition in different environments.

A challenge for the inclusion of floral traits in large-scale evolutionary and ecological studies seems to be their lack of availability in databases. The largest plant database today, TRY, contains limited data about flowers as compared to vegetative and seed traits (Kattge et al., 2020). Although data and floras are increasingly available online, they still represent a small fraction of known plant species, with tropical floras typically being underrepresented (cf Römer et al., 2023). Many previous studies, although they sometimes included many more species, have focused on already available datasets, and are thus often biased towards temperate regions, in particular Europe and the USA (e.g. Díaz et al., 2016; Salguero-Gómez et al., 2016; E-Vojtkó et al., 2022). Other studies have recently compiled data for many species but focus on one or a few traits (e.g. Wang et al., 2021; Wang et al., 2023). One could combine data from different sources to assemble information about multiple traits, but by filtering species based on data completeness, the same geographic and phylogenetic biases are likely to arise. For research questions on angiosperms, data should be assembled based on the representation of angiosperm diversity rather than data availability. This requires combining species-level data from different datasets and different types of literature (scientific studies, floras), an often time-consuming task. In order to efficiently combine more data, we need publicly available data with well-described trait standards, references to the primary literature, and quality checks by botanists who are familiar with the terminology (Sauquet & Magallón, 2018). This would allow to study rare mating or sexual systems (e.g. heterodichogamy, androdioecy) and systems that are not always reported in botanical descriptions (e.g. herkogamy, dichogamy; Cardoso et al., 2018).

The inclusion of floral and pollination-related traits in the description of the plant functional diversity is necessary to improve our understanding of ecosystem functioning. Indeed, vegetative functional diversity has proven instrumental in testing theories of the role of diversity in ecosystems (e.g. Lamanna et al., 2014; Schuldt et al., 2019) but investigating floral diversity in this context may yet yield further insights. Indeed, interactions with pollinators have been identified as playing a role in the maintenance of diversity in plant communities (Wei et al., 2021) and the decrease of pollinator abundance can destabilize the mechanisms promoting species coexistence (Johnson et al., 2022). As we are only starting to standardize floral trait data and make them available, we still have a very incomplete picture about how they influence pollinator diversity and abundance, and the subsequent effects on plant community composition. However, such understanding is urgently needed as pollinators are declining rapidly in many agricultural and semi-natural landscapes (Artamendi et al., 2025), and this angiosperm-wide study provides a framework for future research.

## Acknowledgments

This research was supported by the Fondation pour la Recherche sur la Biodiversité (FRB) through the CESAB project ‘DiveRS’; the FRB, Office Français de la Biodiversité and Ministère de Transition Ecologique project ‘IndicatoRS’; and by the ANR project ‘FloweRS’ (ANR-23-CE02-0003). We thank the DiveRS members Sally Otto and Denis Roze for constructive discussions and feedback on the manuscript. We thank Nicolas Casajus, Nicolas Loiseau and Marion Chartier for advice on data analyses, Solenn Sarton for contributing data, and Maud Calmet for the organisation of the workshops of the DiveRS group. We thank the University of Vienna for hosting the PROTEUS database and eFLOWER server. We thank Susanne Renner and two anonymous reviewers for comments that helped to improve the manuscript.

## Author contributions

SG and JK initiated the project; BA, SB, CB, HdB, MD, SG, JK, MM, JRP, HS, DJS, JS, MV-M, RZ-F and AJH conceived the project; MvB, HS and JS provided data management; MvB, CB, JC, P-AD, SG, JK, MM, DSB, HS and JS entered data; AJH, SG, JK and HS performed analyses; AJH visualized results; AJH, JK and SG drafted the first version of the manuscript; all authors contributed to and approved of the final version.

## Data availability

Data, all scripts, and additional analyses are available at the following github repository: https://github.com/divers-it/rs-traitspace. The original data are also available at data.InDoRES, https://doi.org/10.48579/PRO/HU3QYM

## Supporting Information

**Figure S1:** The amount of missing data per trait after the dataset has been cleaned.

**Figure S2:** The relationships between different pairwise distance matrices.

**Figure S3:** Comparison of the species sampled in the current study and the obtained trait space with species and traits derived from Díaz et al. (2022).

**Figure S4:** Gower’s distances calculated for our data set vs those calculated for a data set derived from Díaz et al., 2022.

**Figure S5:** Comparisons of trait space quality (measured using AUC) among data sets.

**Figure S6:** Simulated vs observed eigenvalues from principal coordinate analyses (PCoA).

**Figure S7:** Third and fourth axes of the principal coordinate analysis on our one-hot encoded data set.

**Figure S8:** Species clustering shown on the first two axes of a principal coordinates analysis (PCoA) on the original data set.

**Figure S9:** Trait values per cluster.

**Figure S10:** The distributions of species along the first two axes of UMAP (uniform manifold approximation) analyses.

**Figure S11:** A Sankey plot depicting flows between clusters as the value of ‘k’ (number of clusters) was sequentially increased.

**Figure S12:** Characterizing robust groups identified using the Partitioning Around Mediods (PAM) clustering approach.

**Figure S13:** Boxplots showing the distribution of values for each quantitative trait in our data set, grouped by reproductive system.

**Table S1:** Plant traits included in this study.

**Table S2:** A list of all species included in the analyses.

**Table S3:** The number of species per order used in this study.

**Table S4:** The correlation of each trait (original data set) with the first four principal coordinate analysis (PCoA) axes.

Notes S1: Trait scoring guide. Instructions for entering trait data in the PROTEUS database.

