## supplementary materials for "An angiosperm-wide perspective on reproductive strategies and floral traits"

#### The following Supporting Information is available for this article:

**Supplementary figures** **2**

**Supplementary tables** **15**

**Notes S1: Trait scoring guide** **27**

**Additional references** **36**

### Supplementary figures

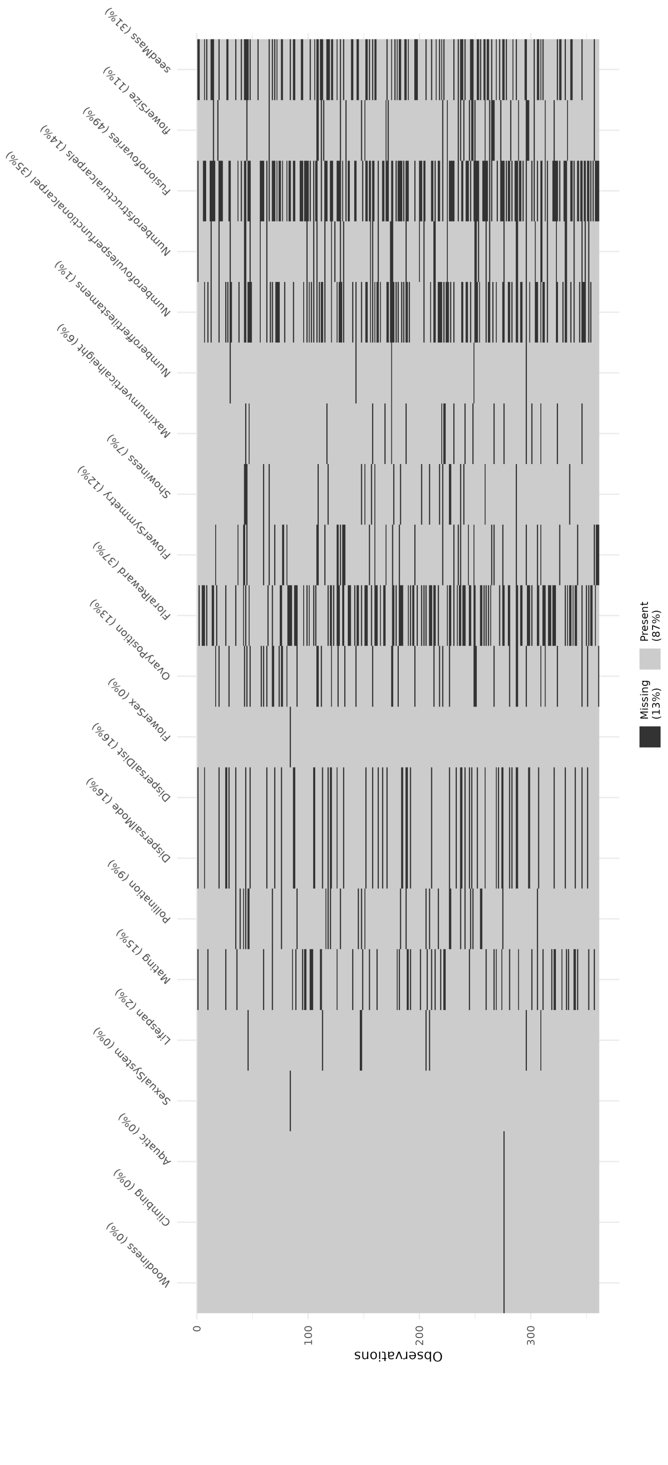


Figure S1: The amount of missing data per trait after the dataset has been cleaned. Traits are in columns and each line is a species in our data set. Black lines in each column indicate where data are missing.

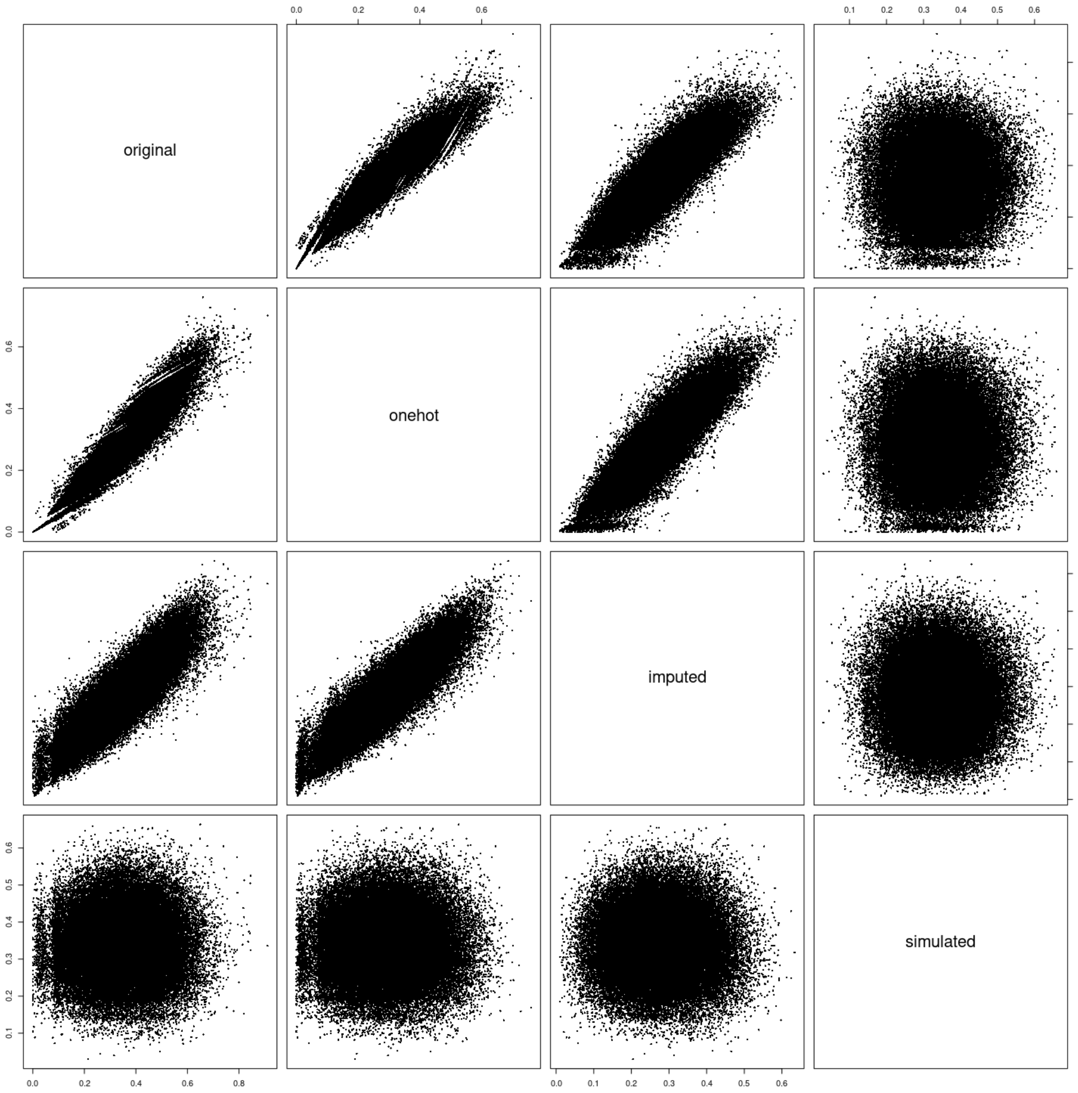


Figure S2: The relationships between different pairwise distance matrices. Along the diagonal are labels indicating each different parwise distance matrix used in this paper, and whether it is on the X or Y axis of each scatterplot. These were made from different data sets, from the top left: the original data set, a one-hot encoded data set, the original data set where missing data were imputed and finally a data set that was simulated using a phylogenetic tree and models of trait evolution (see the methods of the main text for further details).

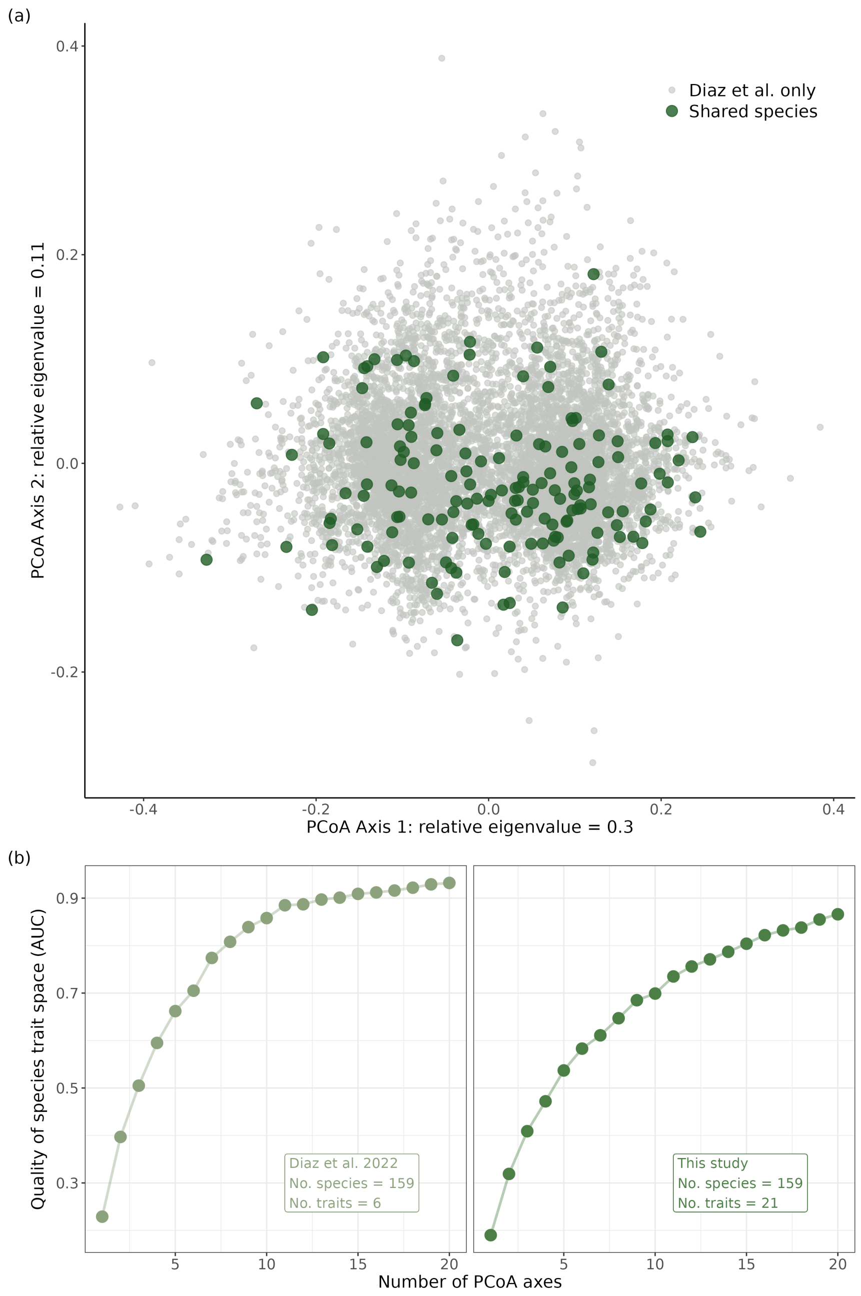


Figure S3: Comparison of the species sampled in the current study and the obtained trait space with species and traits derived from Díaz et al. (2022). Panel (a) shows a scatterplot of a subset of approximately 8,000 species from Díaz et al. (2022) on the first two axes of a principal coordinate analysis (PCoA). Species that overlap with those in this study are highlighted in green. Panel (b) shows two line plots depicting how trait space quality (measured using AUC) changes as PCoA axes are added. Both graphs are made using the same set of 159 species but with trait data from Díaz et al. (2022) (left) and this study (right).

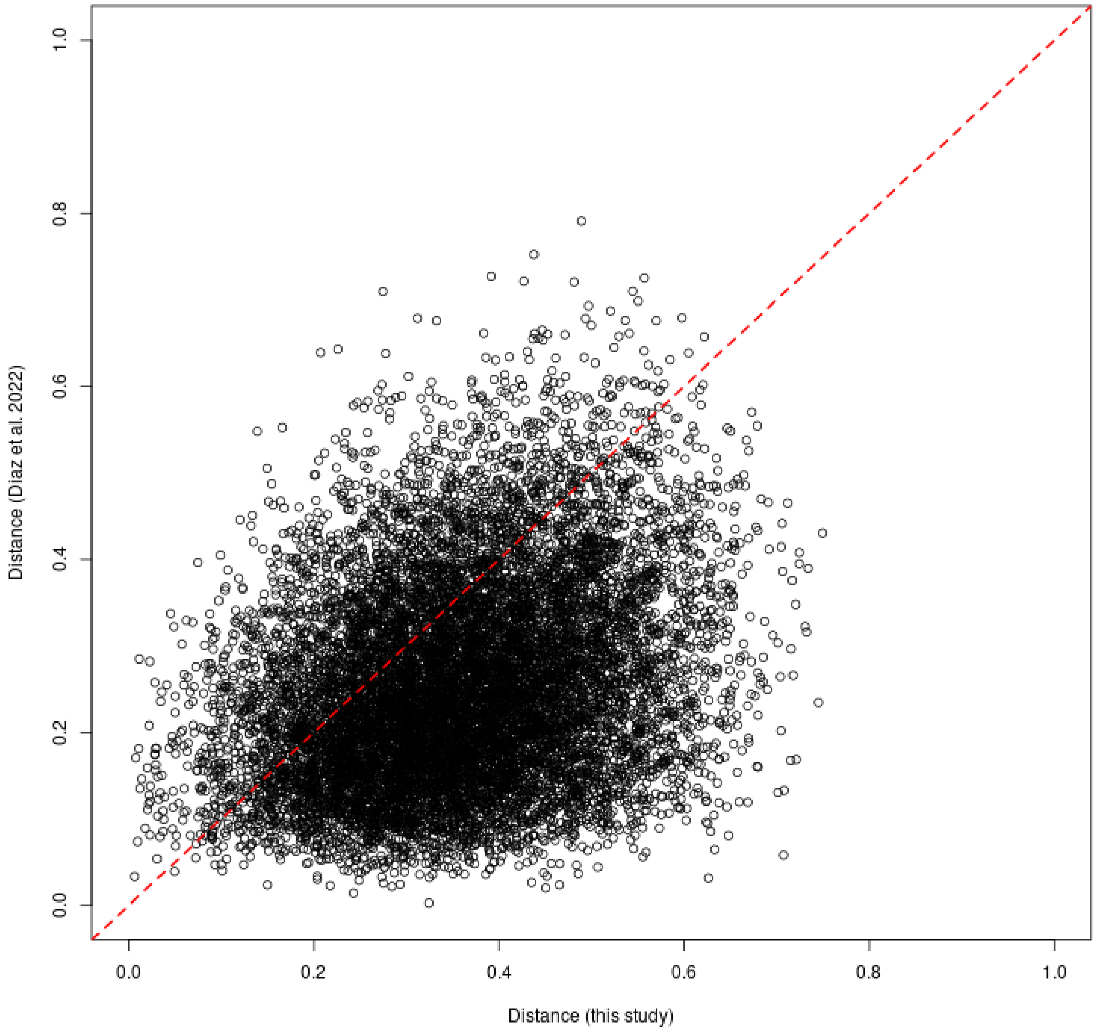


Figure S4: Gower's distances calculated for our data set vs those calculated for a data set derived from Díaz et al., 2022. A total of 159 species were shared across these two data sets and used to make this plot. The dotted red line represents the identity line, which indicates that distances were generally larger in our data set.

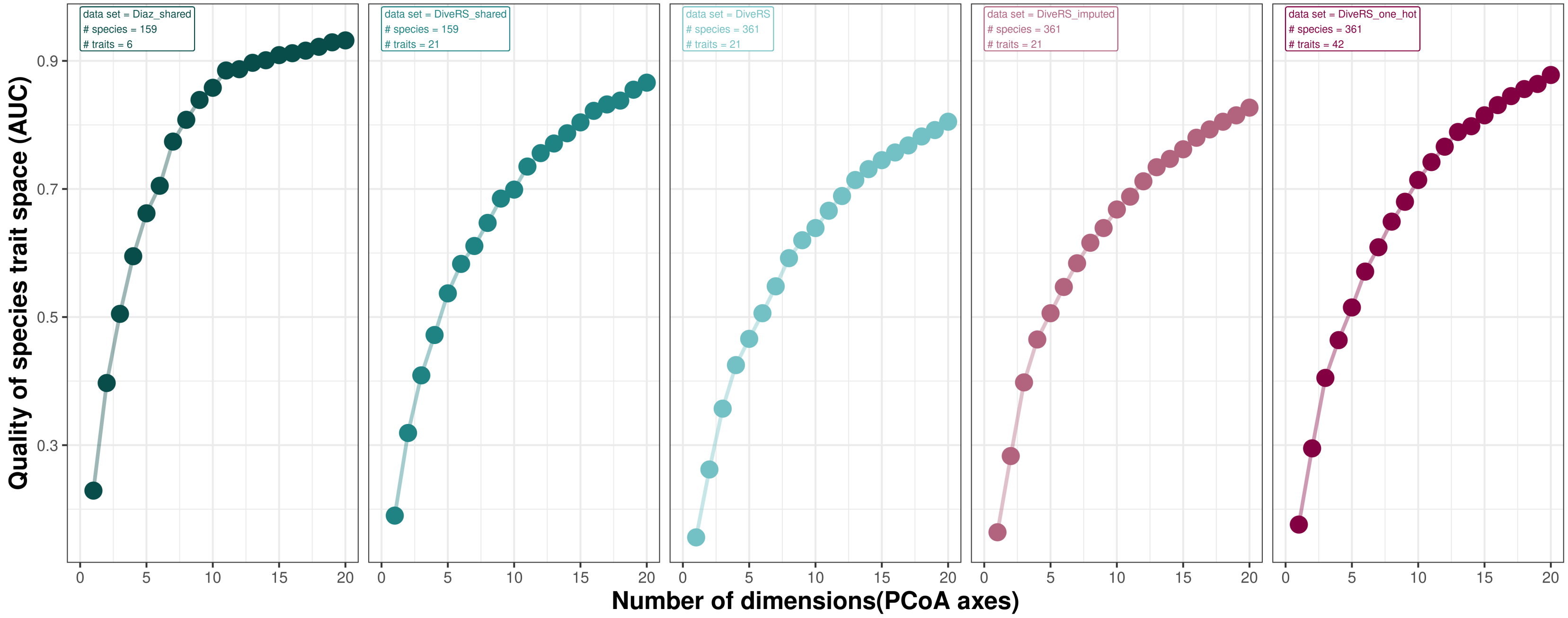


Figure S5: Comparisons of trait space quality (measured using AUC) among data sets. Data sets are indicated in the top left of each panel: 'Diaz shared' contains species present in both Díaz et al., 2022 and our dataset, using the traits of Díaz et al., 2022; 'DiveRS shared': same as previous, but using the traits from the current study; 'DiveRS': original data set used in the current study; 'DiveRS imputed': same as 'DiveRS' but missing data is replaced by imputed data; 'DiveRS one hot': same as 'DiveRS', but using one-hot encoding. Dashed lines and the black point show the number of axes to be kept, as determined by the elbow method, when optimising the trade-off between trait space quality and operationality. See Mouillot et al., 2021 for further details.

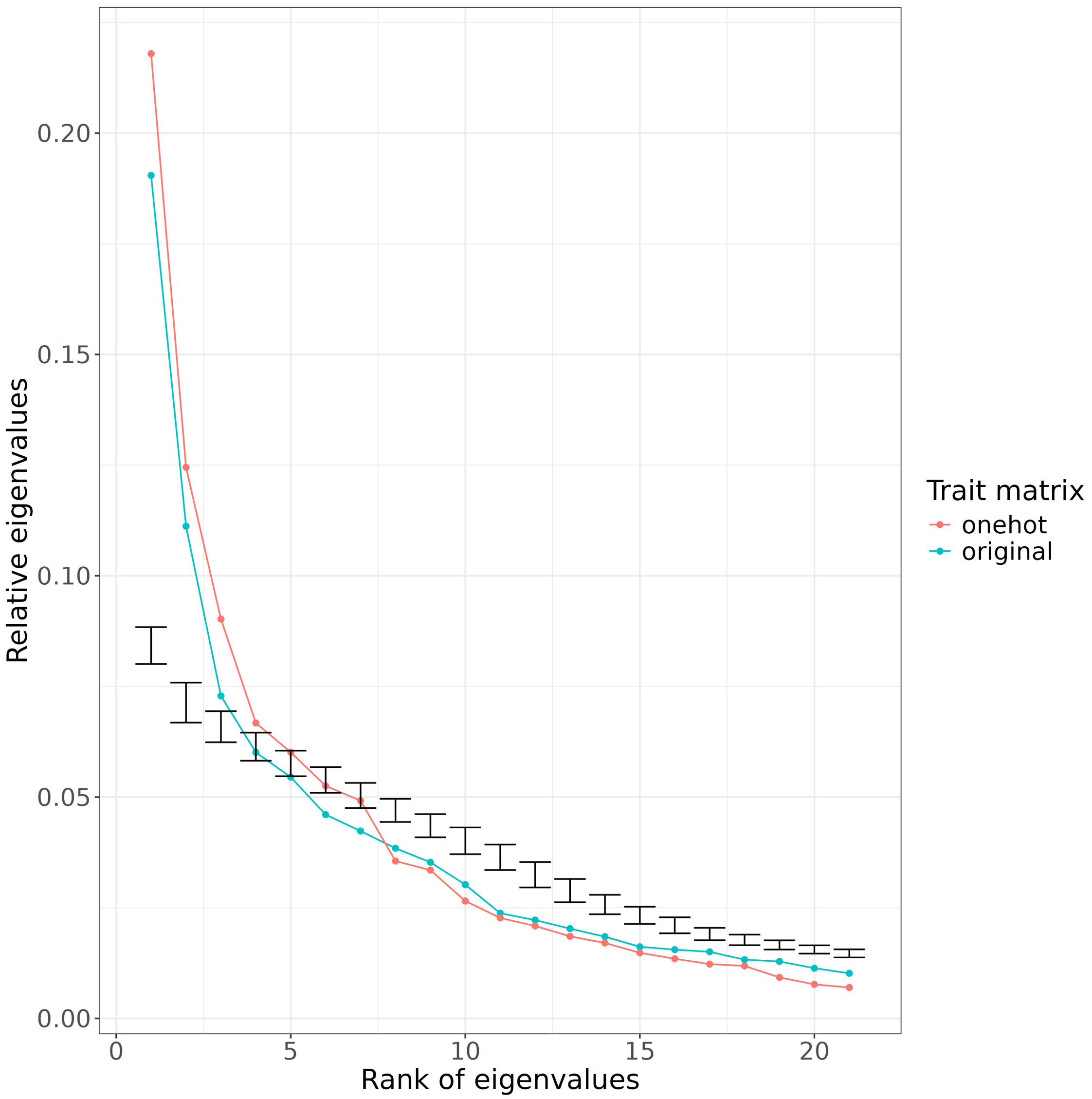


Figure S6: Simulated vs observed eigenvalues from principal cooordinate analyses (PCoA). The black error bars indicate the range of relative eigenvalues for each PCoA axis from 1000 simulated data sets. Blue and red lines show the empirical realtive eigenvalues from our original and one-hot data sets respectively.

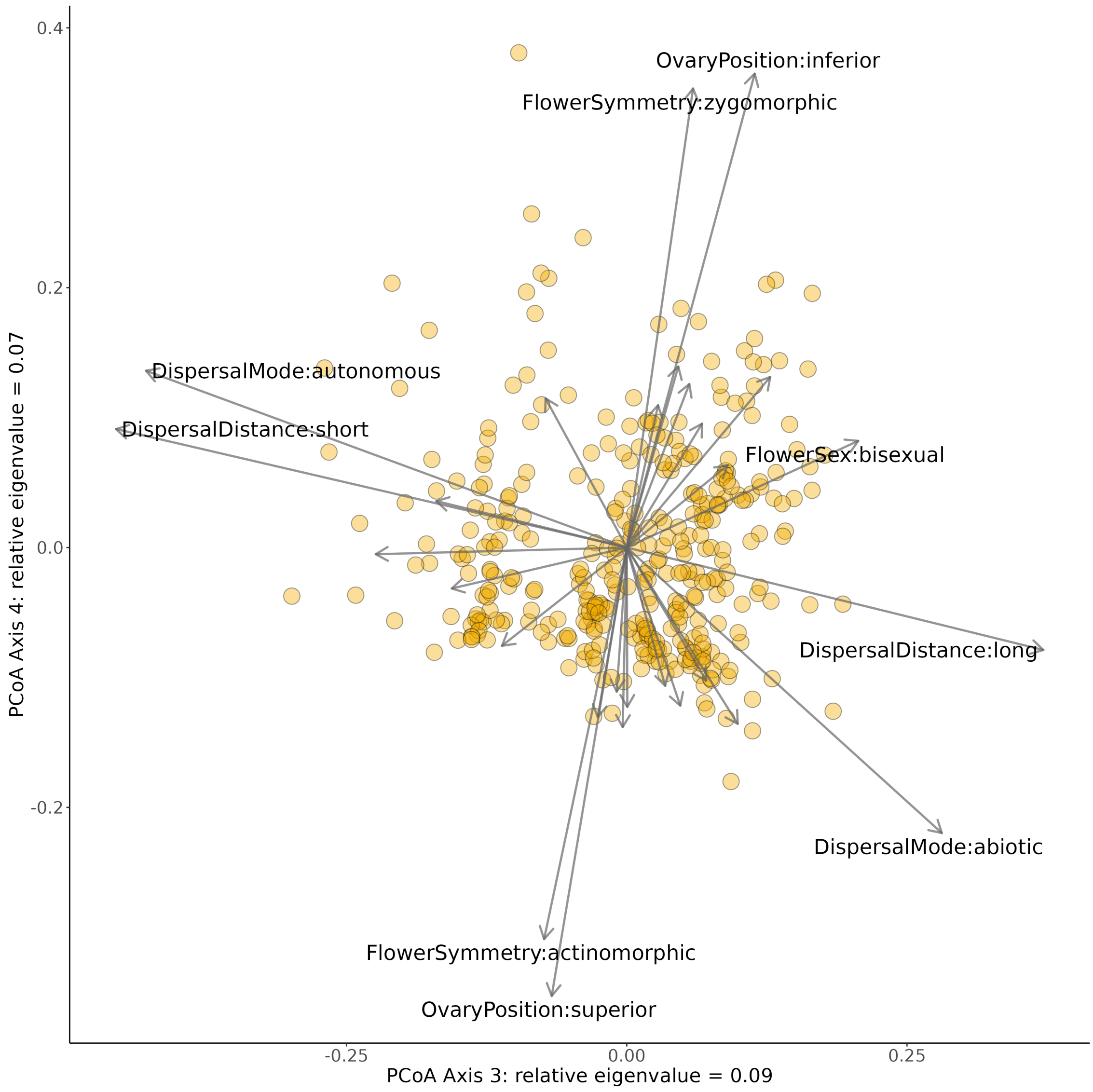


Figure S7: Third and fourth axes of the principal coordinate analysis on our one-hot encoded data set. These axes represent approximately 16% of variation. Using one-hot encoding allowed us to plot arrows indicating how different states and traits affect the trait space, as well as which traits are acting in a similar manner. The length of the arrows indicates the strength of effect of the state and labels were added to arrows representing those states with large effects.

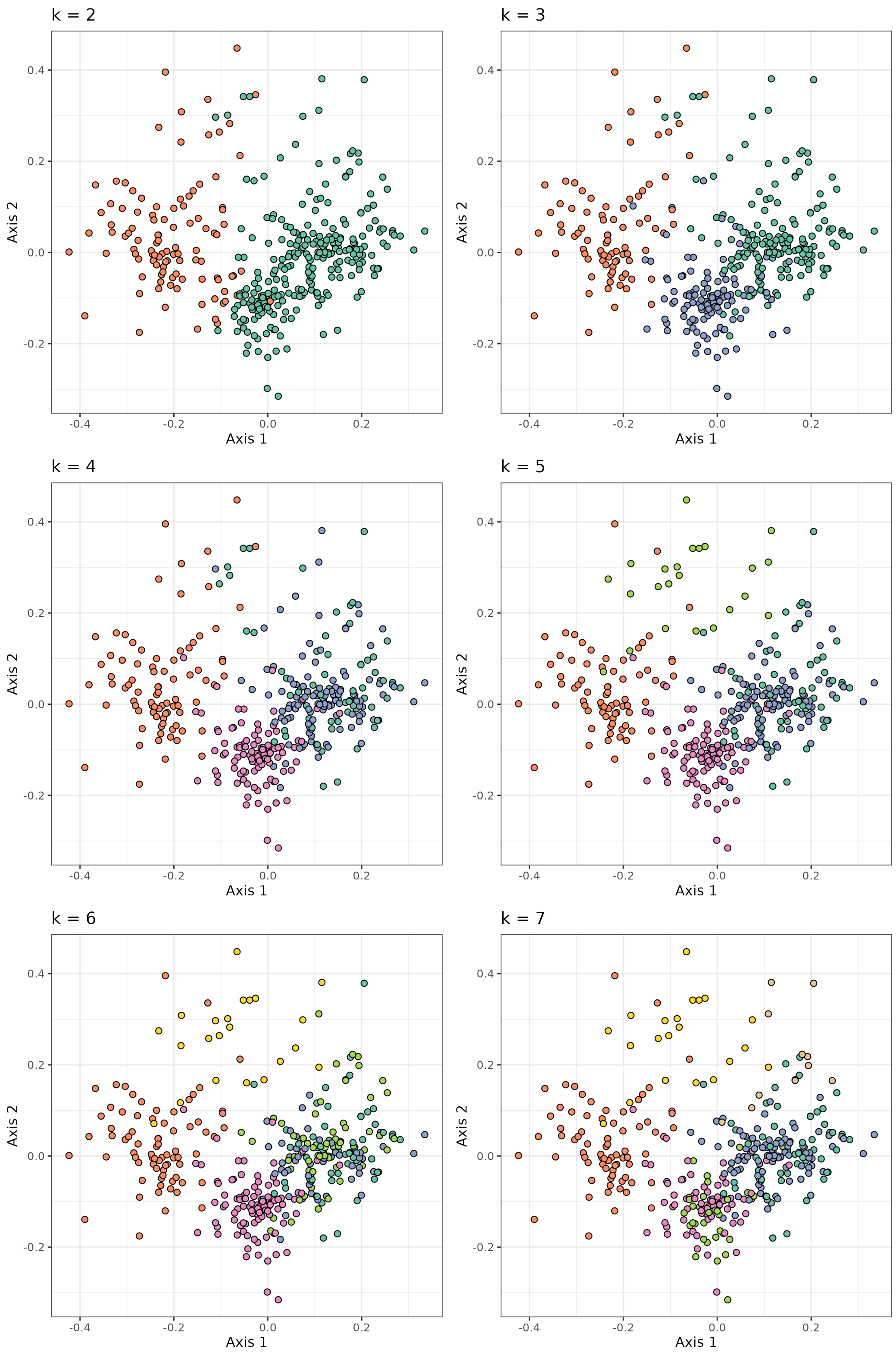


Figure S8: Species clustering shown on the first two axes of a principal coordinates analysis (PCoA) on the original data set. In each scatterplot points are coloured by cluster membership derived from clustering using the partitioning around medoids (PAM) method with increasing values of $k$ (number of clusters), from two to seven clusters.

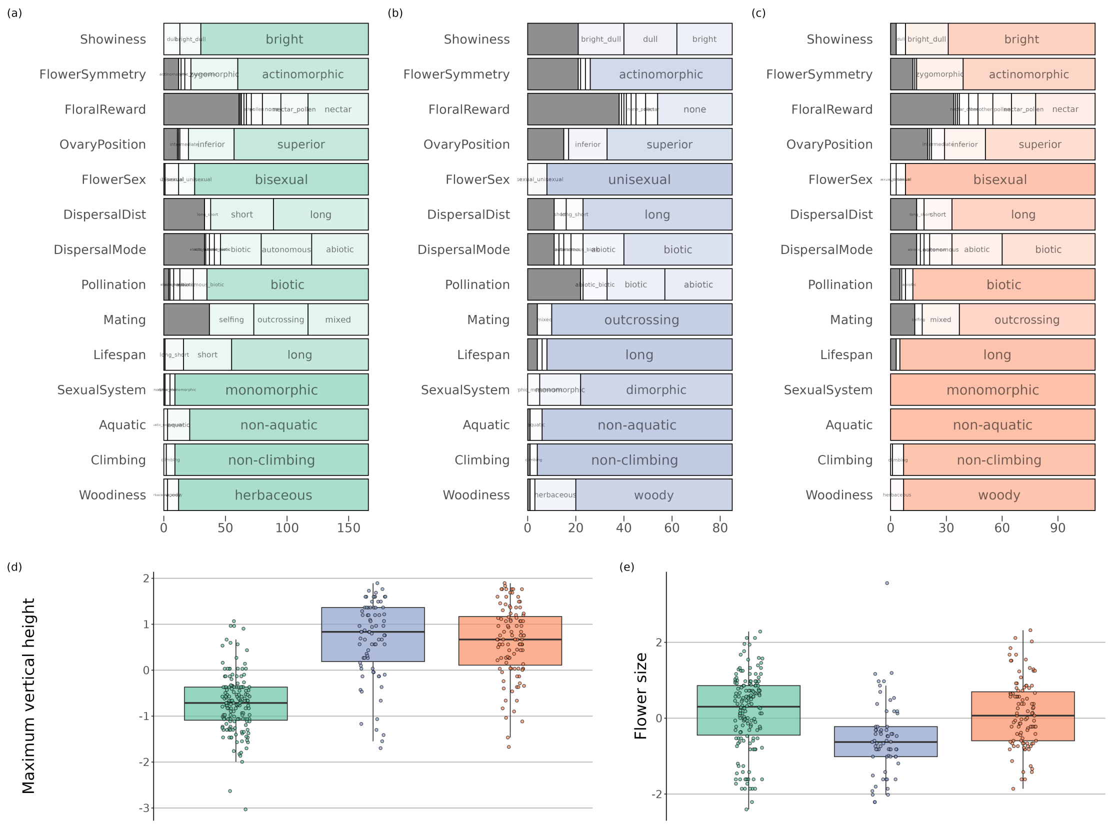


Figure S9: Trait values per cluster. Panels (a-c) show horizontal stacked barplots of the frequency of states of 14 qualitative traits in each cluster from a partitioning around medoids (PAM) analysis when the number of clusters was set to three. On the X-axis is the number of species in the cluster. Dark grey portions of bars indicate missing data. Panels (d) and (e) show the distribution per cluster (after log transformation and scaling) of values for maximum vertical height and flower size respectively.

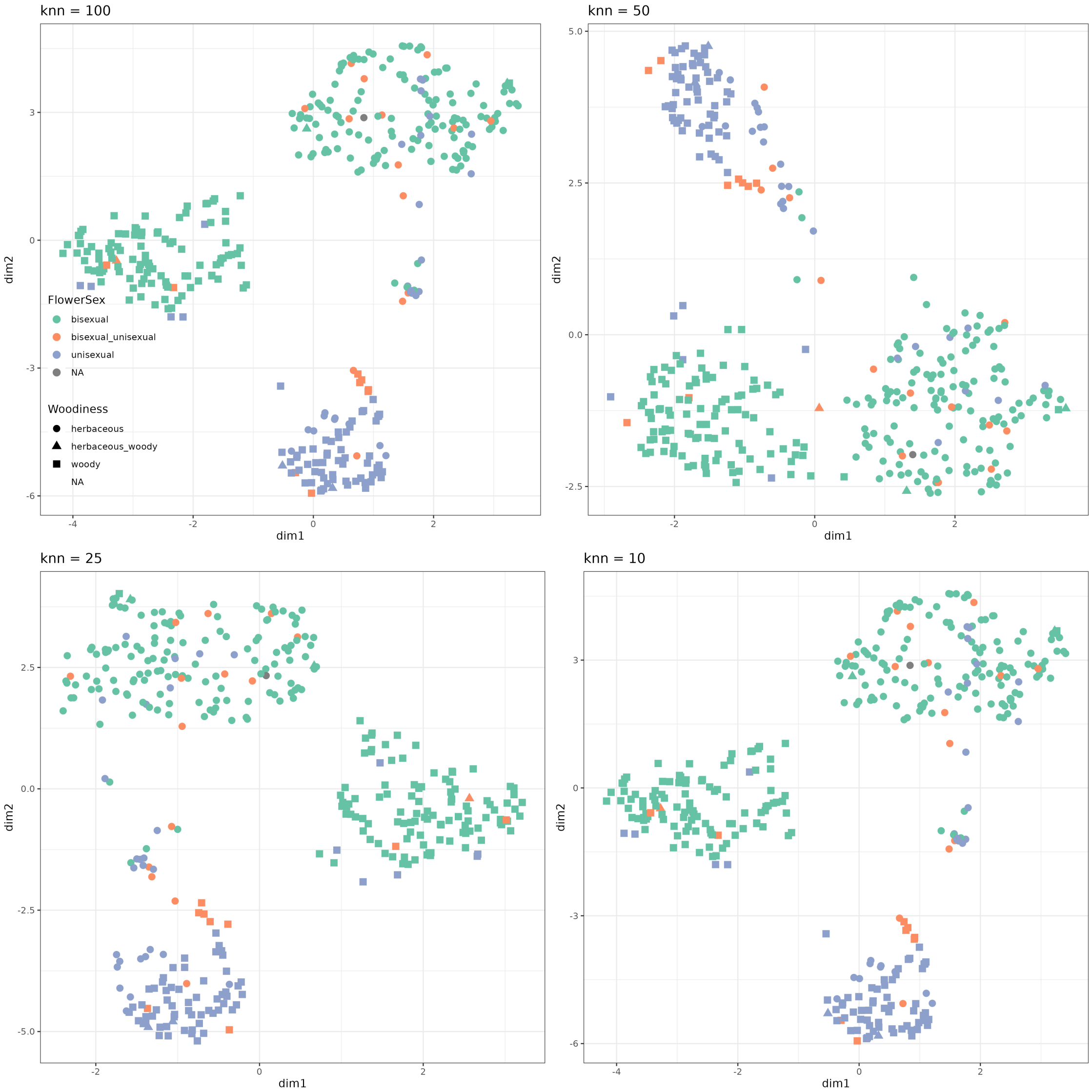


Figure S10: The distributions of species along the first two axes of UMAP (uniform manifold approximation) analyses. Neighbourhood sizes were varied from 100 to 10, shown in the top left of each plot. Larger values allow for a more global view of the manifold and smaller values better preserve local patterns. Points are coloured by flower sex and shapes represent the woodiness trait. As neighbourhood size is decreased three major groups more clearly distinguish themselves.

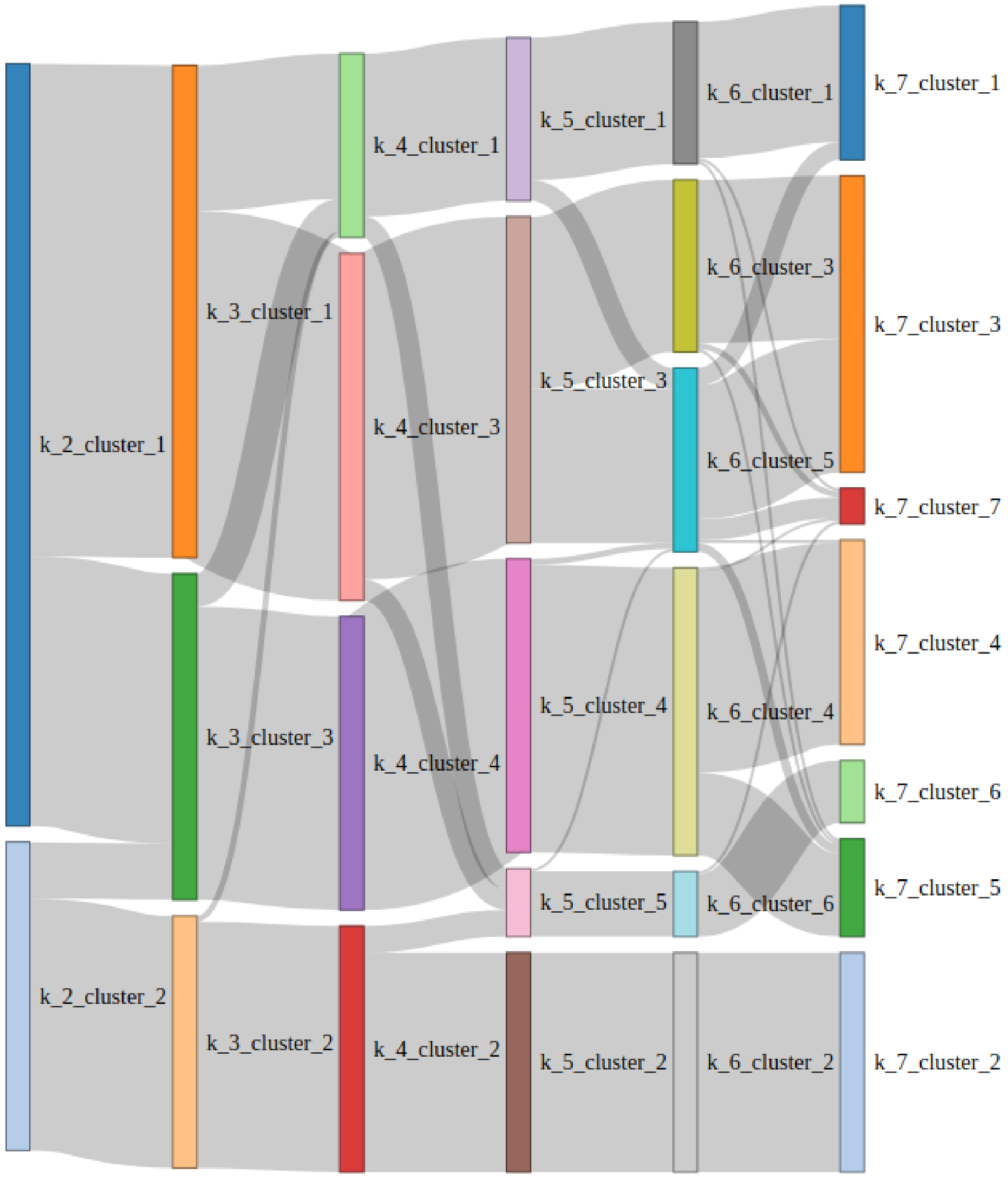


Figure S11: A Sankey plot depicting flows between clusters as the value of 'k' (number of clusters) was sequentially increased. Species were assigned to clusters using paritioning around medoids (PAM) method. Each set of vertical bars is a cluster (from two to seven clusters) and the grey bands between bars show how species moved between clusters when 'k' is changed. Note that individual species are not represented here, merely the number of species that move from one cluster to another (i.e. one species cannot be tracked through all values of 'k').

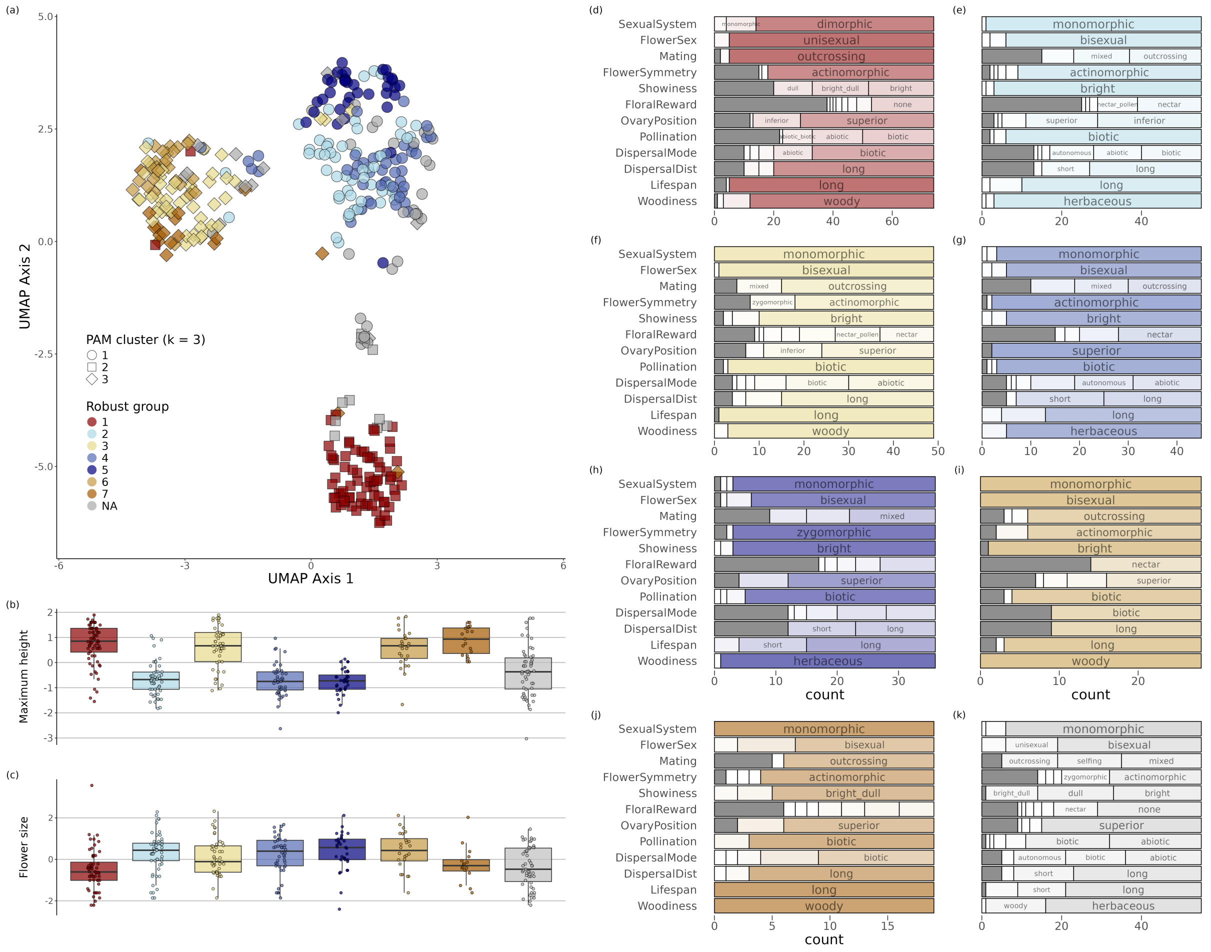


Figure S12: Characterizing robust groups identified using the Partitioning Around Mediods (PAM) clustering approach. Briefly, robust groups are those species that consistently grouped together as the number of clusters was increased (see methods for further details). Seven robust groups were defined, and a final group was made with those species that did not fall into a robust group. Panel (a) shows the distribution of species along the first two axes of a UMAP (uniform manifold approximation) analysis with a neighbourhood size of 10 to represent the final-scale struture in the data. Points are coloured by robust group, and their shape indicates their PAM cluster assignment ($k$ = 3). Boxplots in the bottom left show the distribution (after log transformation and scaling) of values for two traits per cluster: (b) maximum height and (c) flower size. The stacked barplots (panels (d-k)) show the frequency of states for 14 traits for each group. Colours match those in the other panels and bars are split into sections depending on the frequency of each state. Sections representing states with high frequencies are labelled and the dark grey sections correspond to missing data.

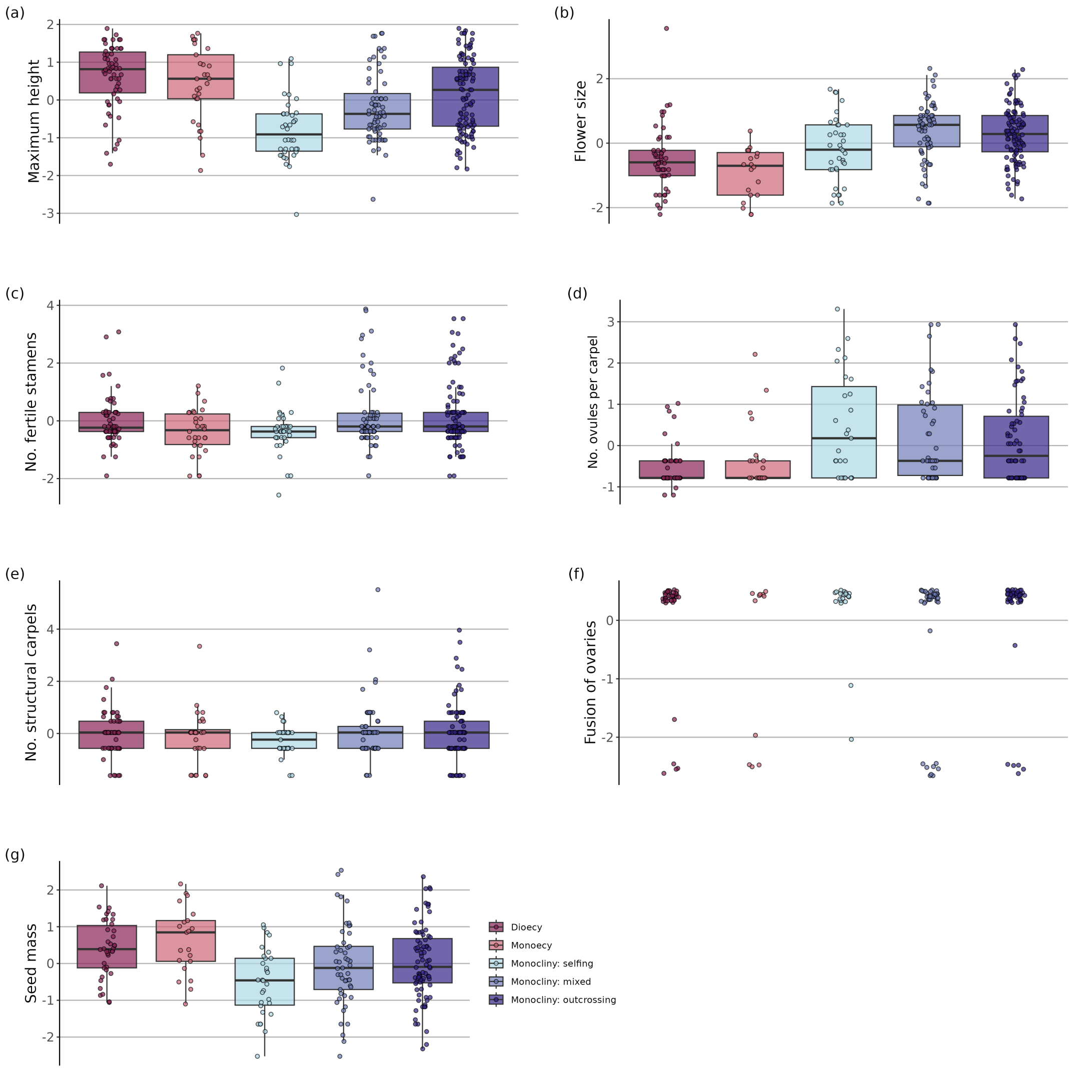


Figure S13: Boxplots showing the distribution of values for each quantitative trait in our data set, grouped by reproductive system. Colours correspond to Figure 4 in the main text.

#### Supplementary tables

Table S1: Plant traits included in this study. The traits actually used in the analyses are presented, i.e. after simplification for categorical traits and removing traits and species with more than 50% missing data. The original trait is the trait recorded in PROTEUS according to the scoring instructions available in Notes S1, except for seed mass. For numerical traits, the ranges (minimum - maximum) are given before transformation, with their units if applicable. Note that averages are used per species so count data (numbers) are not necessarily integers.

| Plant part | Trait | Type (states or ranges [unit]) | Original trait(s) |
|---|---|---|---|
| Entire plant: | Woodiness | categorical (woody, not woody) | Habit |
| vegetative | Aquatic | categorical (aquatic, not aquatic) | Habit |
| | Climbing | categorical (climbing, not climbing) | Habit |
| | Lifespan | categorical (long, short) | Habit, Life history |
| | Maximum vertical height | numerical (0.01 - 50 m) | Maximum vertical height |
| Entire plant: | Sexual system | categorical (monomorphic, dimorphic) | Plant sexual system |
| reproductive | Mating | categorical (selfing, mixed, outcrossing) | Phenotypic mating system, Self-incompatibility system, Outcrossing rate, Plant sexual system[1] |
| Entire flower | Floral reward | categorical (pollen, nectar, other, none) | Floral reward |
| | Flower sex | categorical (unisexual, bisexual) | Floral structural sex |
| | Flower size | numerical (0.08 - 80 cm) | Flower length, Flower diameter[2] |
| | Flower symmetry | categorical (actinomorphic, zygomorphic, other) | Symmetry of perianth |
| | Pollination | categorical (biotic, abiotic, autonomous) | Pollination syndrome |
| | Showiness | categorical (bright, dull) | Main color of perianth at anthesis, Inflorescence attractive organ |

[1]For the original traits 'Self-incompatibility system' and 'Plant sexual system', only the strictly outcrossing states (self-incompatibility and dioecy) were used to determine the mating system

[2]If both were available, the largest dimension was used.

| | | | |
|---|---|---|---|
| | Ovary position | categorical (superior, intermediate, inferior) | Ovary position |
| Female organs | Fusion of ovaries | numerical (0 - 1) | Fusion of ovaries |
| | Number of ovules per carpel | numerical (0.5 - 940) | Number of ovules per functional carpel |
| | Number of structural carpels | numerical (1 - 115) | Number of structural carpels |
| Male organs | Number of fertile stamens | numerical (0.5 - 425) | Number of fertile stamens |
| Dispersal | Dispersal distance | categorical (long, short) | Dispersal syndrome |
| | Dispersal mode | categorical (biotic, abiotic, autonomous) | Dispersal syndrome |
| | Seed mass | numerical (0.001 - 32835 g) | Seed mass |

Table S2: A list of all species included in the analyses.

| Species | Family | Order |
|---|---|---|
| Abolboda pulchella | Xyridaceae | Poales |
| Achatocarpus praecox | Achatocarpaceae | Caryophyllales |
| Acorus calamus | Acoraceae | Acorales |
| Actinidia chinensis | Actinidiaceae | Ericales |
| Aerva javanica | Amaranthaceae | Caryophyllales |
| Aextoxicon punctatum | Aextoxicaceae | Berberidopsidales |
| Agalinis neoscotica | Orobanchaceae | Lamiales |
| Ailanthus altissima | Simaroubaceae | Sapindales |
| Ainsliaea cordifolia | Asteraceae | Asterales |
| Ajuga reptans | Lamiaceae | Lamiales |
| Akebia quinata | Lardizabalaceae | Ranunculales |
| Albizia julibrissin | Fabaceae | Fabales |
| Alepis flavida | Loranthaceae | Santalales |
| Allanblackia floribunda | Clusiaceae | Malpighiales |
| Allium sativum | Amaryllidaceae | Asparagales |
| Allium schoenoprasum | Amaryllidaceae | Asparagales |
| Allocasuarina verticillata | Casuarinaceae | Fagales |
| Alluaudia procera | Didiereaceae | Caryophyllales |
| Alnus alnobetula | Betulaceae | Fagales |
| Aloe thraskii | Asphodelaceae | Asparagales |
| Alphitonia excelsa | Rhamnaceae | Rosales |
| Alseuosmia macrophylla | Alseuosmiaceae | Asterales |
| Alstroemeria aurea | Alstroemeriaceae | Liliales |
| Alyssum simplex | Brassicaceae | Brassicales |
| Amborella trichopoda | Amborellaceae | Amborellales |
| Amsonia tabernaemontana | Apocynaceae | Gentianales |
| Anarthria polyphylla | Restionaceae | Poales |
| Angelonia pubescens | Plantaginaceae | Lamiales |
| Anigozanthos humilis | Haemodoraceae | Commelinales |
| Annona crassiflora | Annonaceae | Magnoliales |
| Antirrhinum majus | Plantaginaceae | Lamiales |
| Apium graveolens | Apiaceae | Apiales |
| Aquilegia canadensis | Ranunculaceae | Ranunculales |
| Arabidopsis thaliana | Brassicaceae | Brassicales |
| Aralidium pinnatifidum | Torricelliaceae | Apiales |
| Arctostaphylos uva-ursi | Ericaceae | Ericales |
| Arum maculatum | Araceae | Alismatales |
| Asparagus officinalis | Asparagaceae | Asparagales |
| Astelia alpina | Asteliaceae | Asparagales |
| Audouinia capitata | Bruniaceae | Bruniales |
| Avicennia marina | Acanthaceae | Lamiales |
| Azima tetracantha | Salvadoraceae | Brassicales |
| Bagassa guianensis | Moraceae | Rosales |
| Balanophora laxiflora | Balanophoraceae | Santalales |
| Balanops pancheri | Balanopaceae | Malpighiales |

| | | |
|---|---|---|
| Bdallophytum americanum | Cytinaceae | Malvales |
| Begonia heracleifolia | Begoniaceae | Cucurbitales |
| Bergeranthus multiceps | Aizoaceae | Caryophyllales |
| Berteroa incana | Brassicaceae | Brassicales |
| Bertholletia excelsa | Lecythidaceae | Ericales |
| Beta vulgaris | Chenopodiaceae | Caryophyllales |
| Bixa orellana | Bixaceae | Malvales |
| Borago officinalis | Boraginaceae | Boraginales |
| Brasenia schreberi | Cabombaceae | Nymphaeales |
| Bretschneidera sinensis | Akaniaceae | Brassicales |
| Bruguiera gymnorhiza | Rhizophoraceae | Malpighiales |
| Buddleja davidii | Scrophulariaceae | Lamiales |
| Bursera simaruba | Burseraceae | Sapindales |
| Butomus umbellatus | Butomaceae | Alismatales |
| Buxus balearica | Buxaceae | Buxales |
| Byrsonima crassifolia | Malpighiaceae | Malpighiales |
| Cabomba caroliniana | Cabombaceae | Nymphaeales |
| Calceolaria purpurea | Calceolariaceae | Lamiales |
| Camellia sinensis | Theaceae | Ericales |
| Canacomyrica monticola | Myricaceae | Fagales |
| Canna indica | Cannaceae | Zingiberales |
| Cannabis sativa | Cannabaceae | Rosales |
| Capparis zeylanica | Capparaceae | Brassicales |
| Cardiocrinum cordatum | Liliaceae | Liliales |
| Carex laxiflora | Cyperaceae | Poales |
| Carica papaya | Caricaceae | Brassicales |
| Caryocar brasiliense | Caryocaraceae | Malpighiales |
| Catalpa speciosa | Bignoniaceae | Lamiales |
| Catharanthus roseus | Apocynaceae | Gentianales |
| Caulophyllum thalictroides | Berberidaceae | Ranunculales |
| Centroplacus glaucinus | Centroplacaceae | Malpighiales |
| Ceratophyllum demersum | Ceratophyllaceae | Ceratophyllales |
| Cercidiphyllum japonicum | Cercidiphyllaceae | Saxifragales |
| Chimaphila umbellata | Ericaceae | Ericales |
| Chimonanthus praecox | Calycanthaceae | Laurales |
| Chloranthus spicatus | Chloranthaceae | Chloranthales |
| Chorispora tenella | Brassicaceae | Brassicales |
| Cinnamomum camphora | Lauraceae | Laurales |
| Cirsium pitcheri | Asteraceae | Asterales |
| Clarkia xantiana | Onagraceae | Myrtales |
| Claytonia virginica | Montiaceae | Caryophyllales |
| Cleistesiopsis bifaria | Orchidaceae | Asparagales |
| Cleomella arborea | Cleomaceae | Brassicales |
| Clethra alnifolia | Clethraceae | Ericales |
| Clutia pulchella | Peraceae | Malpighiales |
| Cobaea scandens | Polemoniaceae | Ericales |
| Cochlospermum vitifolium | Bixaceae | Malvales |
| Coffea arabica | Rubiaceae | Gentianales |

| | | |
|---|---|---|
| Combretum leprosum | Combretaceae | Myrtales |
| Commelina communis | Commelinaceae | Commelinales |
| Cordyline australis | Asparagaceae | Asparagales |
| Coriaria myrtifolia | Coriariaceae | Cucurbitales |
| Coriaria ruscifolia | Coriariaceae | Cucurbitales |
| Cornus florida | Cornaceae | Cornales |
| Corokia cotoneaster | Argophyllaceae | Asterales |
| Corrigiola litoralis | Caryophyllaceae | Caryophyllales |
| Costus woodsonii | Costaceae | Zingiberales |
| Cucurbita pepo | Cucurbitaceae | Cucurbitales |
| Cupaniopsis anacardioides | Sapindaceae | Sapindales |
| Curculigo latifolia | Hypoxidaceae | Asparagales |
| Cuscuta cuspidata | Convolvulaceae | Solanales |
| Cuttsia viburnea | Rousseaceae | Asterales |
| Cyclanthus bipartitus | Cyclanthaceae | Pandanales |
| Cymodocea nodosa | Cymodoceaceae | Alismatales |
| Cypripedium calceolus | Orchidaceae | Asparagales |
| Cyrilla racemiflora | Cyrillaceae | Ericales |
| Dalechampia spathulata | Euphorbiaceae | Malpighiales |
| Dalzellia ceylanica | Podostemaceae | Malpighiales |
| Daphniphyllum macropodum | Daphniphyllaceae | Saxifragales |
| Datisca glomerata | Datiscaceae | Cucurbitales |
| Davidsonia jerseyana | Cunoniaceae | Oxalidales |
| Davilla kunthii | Dilleniaceae | Dilleniales |
| Denhamia celastroides | Celastraceae | Celastrales |
| Diapensia lapponica | Diapensiaceae | Ericales |
| Diospyros australis | Ebenaceae | Ericales |
| Dipterocarpus tempehes | Dipterocarpaceae | Malvales |
| Discaria americana | Rhamnaceae | Rosales |
| Drosophyllum lusitanicum | Drosophyllaceae | Caryophyllales |
| Dryas octopetala | Rosaceae | Rosales |
| Durio zibethinus | Malvaceae | Malvales |
| Eriocaulon parkeri | Eriocaulaceae | Poales |
| Eriogonum arborescens | Polygonaceae | Caryophyllales |
| Eryngium alpinum | Apiaceae | Apiales |
| Erythroxylum laurifolium | Erythroxylaceae | Malpighiales |
| Eschscholzia californica | Papaveraceae | Ranunculales |
| Eucalyptus cladocalyx | Myrtaceae | Myrtales |
| Eupomatia bennettii | Eupomatiaceae | Magnoliales |
| Eupomatia laurina | Eupomatiaceae | Magnoliales |
| Fagus grandifolia | Fagaceae | Fagales |
| Filipendula vulgaris | Rosaceae | Rosales |
| Floerkea proserpinacoides | Limnanthaceae | Brassicales |
| Fumana ericoides | Cistaceae | Malvales |
| Galium aparine | Rubiaceae | Gentianales |
| Gelsemium sempervirens | Gelsemiaceae | Gentianales |
| Gentiana saponaria | Gentianaceae | Gentianales |
| Geranium sanguineum | Geraniaceae | Geraniales |

| | | |
|---|---|---|
| Gerbera jamesonii | Asteraceae | Asterales |
| Gesneria cuneifolia | Gesneriaceae | Lamiales |
| Gomortega keule | Gomortegaceae | Laurales |
| Gonocaryum litorale | Cardiopteridaceae | Cardiopteridales |
| Gossypium hirsutum | Malvaceae | Malvales |
| Grewia occidentalis | Malvaceae | Malvales |
| Griselinia racemosa | Griseliniaceae | Apiales |
| Gyrocarpus americanus | Hernandiaceae | Laurales |
| Hamamelis virginiana | Hamamelidaceae | Saxifragales |
| Hanguana malayana | Hanguanaceae | Commelinales |
| Haumania danckelmaniana | Marantaceae | Zingiberales |
| Hedychium gardnerianum | Zingiberaceae | Zingiberales |
| Helianthus annuus | Asteraceae | Asterales |
| Heliconia irrasa | Heliconiaceae | Zingiberales |
| Herniaria glabra | Caryophyllaceae | Caryophyllales |
| Hevea brasiliensis | Euphorbiaceae | Malpighiales |
| Hirtella physophora | Chrysobalanaceae | Malpighiales |
| Hordeum bulbosum | Poaceae | Poales |
| Hydnocarpus heterophyllus | Achariaceae | Malpighiales |
| Hydrastis canadensis | Ranunculaceae | Ranunculales |
| Hydrocotyle vulgaris | Araliaceae | Apiales |
| Hydrophyllum capitatum | Hydrophyllaceae | Boraginales |
| Hypericum perforatum | Hypericaceae | Malpighiales |
| Ilex opaca | Aquifoliaceae | Aquifoliales |
| Impatiens capensis | Balsaminaceae | Ericales |
| Ipomoea alba | Convolvulaceae | Solanales |
| Iris versicolor | Iridaceae | Asparagales |
| Irvingbaileya australis | Stemonuraceae | Cardiopteridales |
| Juglans mandshurica | Juglandaceae | Fagales |
| Jumellea fragrans | Orchidaceae | Asparagales |
| Juncus effusus | Juncaceae | Poales |
| Kalanchoe daigremontiana | Crassulaceae | Saxifragales |
| Kalmia latifolia | Ericaceae | Ericales |
| Kielmeyera regalis | Calophyllaceae | Malpighiales |
| Kirengeshoma palmata | Hydrangeaceae | Cornales |
| Kirkia wilmsii | Kirkiaceae | Sapindales |
| Lactoris fernandeziana | Lactoridaceae | Piperales |
| Lantana camara | Verbenaceae | Lamiales |
| Lapageria rosea | Smilacaceae | Liliales |
| Larrea tridentata | Zygophyllaceae | Zygophyllales |
| Lemna minor | Araceae | Alismatales |
| Lepidobotrys staudtii | Lepidobotryaceae | Celastrales |
| Limnanthes douglasii | Limnanthaceae | Brassicales |
| Linum perenne | Linaceae | Malpighiales |
| Liparis loeselii | Orchidaceae | Asparagales |
| Liriodendron chinense | Magnoliaceae | Magnoliales |
| Loasa heterophylla | Loasaceae | Cornales |
| Lobelia angulata | Campanulaceae | Asterales |

| | | |
|---|---|---|
| Ludwigia adscendens | Onagraceae | Myrtales |
| Lythrum salicaria | Lythraceae | Myrtales |
| Malesherbia linearifolia | Passifloraceae | Malpighiales |
| Malpighia emarginata | Malpighiaceae | Malpighiales |
| Manihot esculenta | Euphorbiaceae | Malpighiales |
| Manilkara zapota | Sapotaceae | Ericales |
| Mappia nimmoniana | Icacinaceae | Icacinales |
| Marathrum rubrum | Podostemaceae | Malpighiales |
| Mayaca fluviatilis | Mayacaceae | Poales |
| Mazus pumilus | Mazaceae | Lamiales |
| Medicago sativa | Fabaceae | Fabales |
| Melocactus curvispinus | Cactaceae | Caryophyllales |
| Menispermum canadense | Menispermaceae | Ranunculales |
| Menyanthes trifoliata | Menyanthaceae | Asterales |
| Miconia affinis | Melastomataceae | Myrtales |
| Mirabilis jalapa | Nyctaginaceae | Caryophyllales |
| Misodendrum linearifolium | Misodendraceae | Santalales |
| Mollugo verticillata | Molluginaceae | Caryophyllales |
| Monotropa hypopitys | Ericaceae | Ericales |
| Montinia caryophyllacea | Montiniaceae | Solanales |
| Moringa oleifera | Moringaceae | Brassicales |
| Myriophyllum sibiricum | Haloragaceae | Saxifragales |
| Myristica fragrans | Myristicaceae | Magnoliales |
| Myrtus communis | Myrtaceae | Myrtales |
| Narthecium ossifragum | Nartheciaceae | Dioscoreales |
| Nelumbo lutea | Nelumbonaceae | Proteales |
| Nelumbo nucifera | Nelumbonaceae | Proteales |
| Nemesia strumosa | Scrophulariaceae | Lamiales |
| Nepenthes macfarlanei | Nepenthaceae | Caryophyllales |
| Nicotiana tabacum | Solanaceae | Solanales |
| Nolana humifusa | Solanaceae | Solanales |
| Nothofagus dombeyi | Nothofagaceae | Fagales |
| Nuphar advena | Nymphaeaceae | Nymphaeales |
| Nymphaea odorata | Nymphaeaceae | Nymphaeales |
| Nymphoides peltata | Menyanthaceae | Asterales |
| Nyssa ogeche | Nyssaceae | Cornales |
| Ochroma pyramidale | Malvaceae | Malvales |
| Octoknema borealis | Octoknemaceae | Santalales |
| Octomeles sumatrana | Tetramelaceae | Cucurbitales |
| Olea europaea | Oleaceae | Lamiales |
| Oryza sativa | Poaceae | Poales |
| Ottelia acuminata | Hydrocharitaceae | Alismatales |
| Oxalis corniculata | Oxalidaceae | Oxalidales |
| Paeonia californica | Paeoniaceae | Saxifragales |
| Panax quinquefolius | Araliaceae | Apiales |
| Pandanus tectorius | Pandanaceae | Pandanales |
| Pennantia corymbosa | Pennantiaceae | Apiales |
| Perrottetia ovata | Dipentodontaceae | Huerteales |

| | | |
|---|---|---|
| Petrosavia sakuraii | Petrosaviaceae | Petrosaviales |
| Petunia axillaris | Solanaceae | Solanales |
| Peumus boldus | Monimiaceae | Laurales |
| Phelline comosa | Phellinaceae | Asterales |
| Phoenix dactylifera | Arecaceae | Arecales |
| Phormium tenax | Asphodelaceae | Asparagales |
| Phragmites australis | Poaceae | Poales |
| Phryma leptostachya | Phrymaceae | Lamiales |
| Phyllanthus niruri | Phyllanthaceae | Malpighiales |
| Physena madagascariensis | Physenaceae | Caryophyllales |
| Picramnia polyantha | Picramniaceae | Picramniales |
| Pinguicula alpina | Lentibulariaceae | Lamiales |
| Piper amalago | Piperaceae | Piperales |
| Piper arboreum | Piperaceae | Piperales |
| Pitcairnia albiflos | Bromeliaceae | Poales |
| Pittosporum dasycaulon | Pittosporaceae | Apiales |
| Plantago lanceolata | Plantaginaceae | Lamiales |
| Platea latifolia | Metteniusaceae | Metteniusales |
| Plocosperma buxifolium | Plocospermataceae | Lamiales |
| Plumbago auriculata | Plumbaginaceae | Caryophyllales |
| Podophyllum peltatum | Berberidaceae | Ranunculales |
| Poeppigia procera | Fabaceae | Fabales |
| Polycarpon tetraphyllum | Caryophyllaceae | Caryophyllales |
| Polygala lewtonii | Polygalaceae | Fabales |
| Pontederia crassipes | Pontederiaceae | Commelinales |
| Populus alba | Salicaceae | Malpighiales |
| Portulaca oleracea | Portulacaceae | Caryophyllales |
| Posidonia australis | Posidoniaceae | Alismatales |
| Posoqueria latifolia | Rubiaceae | Gentianales |
| Potamogeton berchtoldii | Potamogetonaceae | Alismatales |
| Primula sieboldii | Primulaceae | Ericales |
| Prionotes cerinthoides | Ericaceae | Ericales |
| Prunus persica | Rosaceae | Rosales |
| Putranjiva roxburghii | Putranjivaceae | Malpighiales |
| Qualea grandiflora | Vochysiaceae | Myrtales |
| Quercus rubra | Fagaceae | Fagales |
| Quinchamalium chilense | Schoepfiaceae | Santalales |
| Quintinia hyehenensis | Paracryphiaceae | Paracryphiales |
| Rafflesia keithii | Rafflesiaceae | Malpighiales |
| Ranunculus acris | Ranunculaceae | Ranunculales |
| Ranunculus ficaria | Ranunculaceae | Ranunculales |
| Ravenala madagascariensis | Strelitziaceae | Zingiberales |
| Reseda lutea | Resedaceae | Brassicales |
| Rhexia virginica | Melastomataceae | Myrtales |
| Rhizophora stylosa | Rhizophoraceae | Malpighiales |
| Ribes aureum | Grossulariaceae | Saxifragales |
| Roridula dentata | Roridulaceae | Ericales |
| Roridula gorgonias | Roridulaceae | Ericales |

| | | |
|---|---|---|
| Roupala montana | Proteaceae | Proteales |
| Rourea induta | Connaraceae | Oxalidales |
| Ruellia ciliatiflora | Acanthaceae | Lamiales |
| Ruta graveolens | Rutaceae | Sapindales |
| Sagina procumbens | Caryophyllaceae | Caryophyllales |
| Sagittaria latifolia | Alismataceae | Alismatales |
| Salpiglossis sinuata | Solanaceae | Solanales |
| Santalum spicatum | Santalaceae | Santalales |
| Sarracenia purpurea | Sarraceniaceae | Ericales |
| Saxifraga cernua | Saxifragaceae | Saxifragales |
| Scaevola montana | Goodeniaceae | Asterales |
| Schinus molle | Anacardiaceae | Sapindales |
| Schisandra chinensis | Schisandraceae | Austrobaileyales |
| Schizanthus grahamii | Solanaceae | Solanales |
| Scleranthus annuus | Caryophyllaceae | Caryophyllales |
| Senecio vulgaris | Asteraceae | Asterales |
| Sesamum indicum | Pedaliaceae | Lamiales |
| Sextonia rubra | Lauraceae | Laurales |
| Silene vulgaris | Caryophyllaceae | Caryophyllales |
| Simmondsia chinensis | Simmondsiaceae | Caryophyllales |
| Sinia rhodoleuca | Ochnaceae | Malpighiales |
| Sinojackia xylocarpa | Styracaceae | Ericales |
| Solanum dulcamara | Solanaceae | Solanales |
| Souroubea guianensis | Marcgraviaceae | Ericales |
| Spergularia marina | Caryophyllaceae | Caryophyllales |
| Spigelia marilandica | Loganiaceae | Gentianales |
| Spiranthes cernua | Orchidaceae | Asparagales |
| Spondias tuberosa | Anacardiaceae | Sapindales |
| Staphylea trifolia | Staphyleaceae | Crossosomatales |
| Stellaria media | Caryophyllaceae | Caryophyllales |
| Sterculia apetala | Malvaceae | Malvales |
| Stylidium graminifolium | Stylidiaceae | Asterales |
| Swietenia macrophylla | Meliaceae | Sapindales |
| Syringa vulgaris | Oleaceae | Lamiales |
| Tacca chantrieri | Taccaceae | Dioscoreales |
| Tacca leontopetaloides | Taccaceae | Dioscoreales |
| Talinum paniculatum | Talinaceae | Caryophyllales |
| Tectona grandis | Lamiaceae | Lamiales |
| Ternstroemia crassifolia | Pentaphylacaceae | Ericales |
| Tetracentron sinense | Trochodendraceae | Trochodendrales |
| Tetratheca aphylla | Elaeocarpaceae | Oxalidales |
| Tetroncium magellanicum | Juncaginaceae | Alismatales |
| Thalictrum minus | Ranunculaceae | Ranunculales |
| Theobroma cacao | Malvaceae | Malvales |
| Thymelaea hirsuta | Thymelaeaceae | Malvales |
| Titanotrichum oldhamii | Gesneriaceae | Lamiales |
| Trachelium caeruleum | Campanulaceae | Asterales |
| Tragopogon dubius | Asteraceae | Asterales |

| | | |
|---|---|---|
| Tribulus terrestris | Zygophyllaceae | Zygophyllales |
| Trichilia emetica | Meliaceae | Sapindales |
| Trillium erectum | Melanthiaceae | Liliales |
| Trithuria submersa | Hydatellaceae | Nymphaeales |
| Trochocarpa laurina | Ericaceae | Ericales |
| Tropaeolum majus | Tropaeolaceae | Brassicales |
| Tropaeolum tricolor | Tropaeolaceae | Brassicales |
| Turnera ulmifolia | Passifloraceae | Malpighiales |
| Typha latifolia | Typhaceae | Poales |
| Urtica dioica | Urticaceae | Rosales |
| Vaccinium uliginosum | Ericaceae | Ericales |
| Valeriana officinalis | Caprifoliaceae | Dipsacales |
| Vellozia squamata | Velloziaceae | Pandanales |
| Verbascum thapsus | Scrophulariaceae | Lamiales |
| Veronica anagallis-aquatica | Plantaginaceae | Lamiales |
| Viburnum rufidulum | Adoxaceae | Dipsacales |
| Viola pubescens | Violaceae | Malpighiales |
| Vitex negundo | Lamiaceae | Lamiales |
| Vitis labrusca | Vitaceae | Vitales |
| Viviania marifolia | Francoaceae | Geraniales |
| Vriesea carinata | Bromeliaceae | Poales |
| Warburgia ugandensis | Canellaceae | Canellales |
| Wurmbea biglandulosa | Colchicaceae | Liliales |
| Xanthosoma sagittifolium | Araceae | Alismatales |
| Ximenia americana | Ximeniaceae | Santalales |
| Zea mays | Poaceae | Poales |
| Zelkova serrata | Ulmaceae | Rosales |
| Zostera marina | Zosteraceae | Alismatales |

Table S3: The number of species per order used in this study.

| Order | Frequency | Order | Frequency |
|---|---|---|---|
| Malpighiales | 28 | Dioscoreales | 3 |
| Caryophyllales | 25 | Pandanales | 3 |
| Lamiales | 25 | Piperales | 3 |
| Ericales | 23 | Proteales | 3 |
| Asterales | 16 | Boraginales | 2 |
| Brassicales | 15 | Cardiopteridales | 2 |
| Asparagales | 14 | Celastrales | 2 |
| Poales | 13 | Dipsacales | 2 |
| Malvales | 12 | Geraniales | 2 |
| Alismatales | 11 | Zygophyllales | 2 |
| Ranunculales | 10 | Acorales | 1 |
| Myrtales | 9 | Amborellales | 1 |
| Rosales | 9 | Aquifoliales | 1 |
| Sapindales | 9 | Arecales | 1 |
| Solanales | 9 | Austrobaileyales | 1 |
| Apiales | 8 | Berberidopsidales | 1 |
| Gentianales | 8 | Bruniales | 1 |
| Saxifragales | 8 | Buxales | 1 |
| Fagales | 7 | Canellales | 1 |
| Santalales | 7 | Ceratophyllales | 1 |
| Cucurbitales | 6 | Chloranthales | 1 |
| Laurales | 6 | Crossosomatales | 1 |
| Zingiberales | 6 | Dilleniales | 1 |
| Liliales | 5 | Huerteales | 1 |
| Magnoliales | 5 | Icacinales | 1 |
| Nymphaeales | 5 | Metteniusales | 1 |
| Commelinales | 4 | Paracryphiales | 1 |
| Cornales | 4 | Petrosaviales | 1 |
| Fabales | 4 | Picramniales | 1 |
| Oxalidales | 4 | Trochodendrales | 1 |
| | | Vitales | 1 |

Table S4: The correlation of each trait (original data set) with the first four principal coordinate analysis (PCoA) axes. Correlations of quantitative traits are represented by Pearson's coefficient and those of qualitative traits were estimated using analysis of variance (ANOVA).

| Trait | Axis.1 | Axis.2 | Axis.3 | Axis.4 |
|---|---|---|---|---|
| Aquatic | 0.012 | 0.143 | 0.06 | 0.031 |
| Climbing | 0.004 | 0.021 | 0.009 | 0.007 |
| Dispersal distance | 0.22 | 0.027 | 0.625 | 0.015 |
| Dispersal mode | 0.263 | 0.108 | 0.656 | 0.047 |
| Floral reward | 0.361 | 0.507 | 0.122 | 0.034 |
| Flower sex | 0.574 | 0.132 | 0.039 | 0.078 |
| Flower size | 0.27 | -0.395 | 0.016 | -0.12 |
| Flower symmetry | 0.149 | 0.017 | 0.028 | 0.32 |
| Fusion of ovaries | 0.029 | -0.149 | -0.046 | -0.095 |
| Lifespan | 0.203 | 0.131 | 0 | 0.005 |
| Mating | 0.346 | 0.099 | 0.053 | 0.061 |
| Maximum height | -0.618 | -0.383 | 0.098 | 0.157 |
| No. fertile stamens | -0.061 | -0.349 | -0.045 | 0.216 |
| No. ovules per carpel | 0.383 | -0.221 | -0.059 | 0.035 |
| No. structural carpels | 0.033 | -0.202 | -0.065 | 0.12 |
| Ovary position | 0.009 | 0.066 | 0.177 | 0.302 |
| Pollination | 0.272 | 0.524 | 0.08 | 0.05 |
| Seed mass | -0.458 | -0.29 | 0.024 | 0.247 |
| Sexual system | 0.482 | 0.046 | 0.072 | 0.087 |
| Showiness | 0.184 | 0.318 | 0.049 | 0.139 |
| Woodiness | 0.416 | 0.279 | 0.007 | 0.064 |

### Notes S1: Trait scoring guide

#### General instructions:

- Don't score if you don't have a reference or an observation for the character
- Prefer data from wild populations, or at least wild genotypes (e.g. plants of wild origin cultivated in botanical gardens or greenhouses). If only data from cultivated crop or ornamental populations are available, please indicate it as a note
- If there are multiple states within the species (polymorphism, subspecies, varieties) code these as separate values / states. There is currently no way to link polymorphism between several traits, so if female flowers are large and blue and male flowers small and green, the only way to indicate this linkage is through comments.
- For rare traits, information is only available for the species possessing them. Please do not score such traits as "absent" yourself if you can't find any information, but only if there is an explicit mention of the absence in the species you're scoring.
- Many traits can be scored from general botanical literature (floras), but some traits (e.g. outcrossing traits, pollination, pollen) require specific literature.

#### Plant-level traits

##### 1. Habit (D1)

Tree, shrub, liana, herb, vine, aquatic herb, subterranean. Lianas are woody climbers (including "climbing shrubs" etc), vines herbaceous climbers (thus, *Vitis vinifera* is a liana). Trees are usually taller than shrubs, and are characterized by one or several trunks, while shrubs have many ramifications. Aquatic herbs are plants that have their leaves at (e.g. *Nymphaea*) or under the water surface (e.g. *Ceratophyllum*), or are free-floating (e.g. *Eichhornia*).

##### 2. Maximum vertical height (C1)

Maximum vertical height (i.e. not the height at first flowering). Indicate in m. Can be a single value or a range of values.

##### 4. Plant sexual system (D1)

This trait combines flower sex and the distribution of the flowers among individual plants. In bisexuality and monoecy (including gynomonoecy and andromonoecy), plants are hermaphroditic, but the distinction is made at the flower level.

**Bisexual:** all individuals have perfect (i.e., bisexual) flowers; some authors use cosexuality or hermaphroditism for this state.

**Monoecious:** all individuals have both male and female flowers

**Dioecious:** each individual is either completely male or completely female.

**Gynomonoecious:** all individuals have both female and perfect flowers

**Gynodioecious:** each individual is either completely female or completely hermaphrodite. Hermaphrodites can be of two kinds, depending on the species (if you have the information, please indicate as a comment). Either each hermaphrodite individual has perfect flowers, or is monoecious (or gyno- or andromonoecious).

**Andromonoecious:** all individuals have both male and perfect flowers

**Androdioecious:** each individual is either completely male or completely hermaphrodite. Hermaphrodites can be of two kinds, depending on the species (if you have the information, please indicate as a comment). Either each hermaphrodite individual has perfect flowers, or is monoecious (or gyno- or andromonoecious).

**Trimonoecy:** all individuals have male, female and perfect flowers.

**Polygamous:** everything that doesn't fit in the preceding categories. Leave a comment.

##### 5. Life history (D1)

Annual, biennial, perennial monocarpic, perennial polycarpic, perennial (undetermined). If possible, for perennial species, indicate whether it flowers only once and then dies (monocarpic), or whether it can flower multiple times (polycarpic). If the information is not available, choose "perennial (undetermined)". If the plant can have several lifecycles a year, it should be considered annual (leave a comment).

##### 80. Phenotypic mating system (D1)

Score this trait only if the mating system is explicitly characterized and noted in a study. Please do not try to infer selfing or outcrossing yourself from flower showiness! Studies might use a combination of floral characters and/or empirical tests (ex P/O ratio, dichogamy, seedset from bagged and unbagged plants,...) in a comparative way, i.e. several species have been characterized in the same study. Use the categorization given by the author. If they used more than three categories, group them into three:

**predominantly selfing:** selfing, predominantly selfing, autogamy, obligate autogamy, facultative autogamy

**mixed:** mixed mating, facultative xenogamy

**predominantly outcrossing:** outcrossing, predominantly outcrossing, obligate outcrossing, xenogamy, obligate xenogamy

If the study provides information such as P/O ratio, autonomous selfing, outcrossing rate, etc.. but does not categorize or qualify the mating system don't do it yourself. We here also score apomixis, which is not a form of selfing strictly speaking, but is predicted to have some similar evolutionary, ecological and genomic consequences.

##### 83. Self-incompatibility system (genetic) (D1)

The basic scoring is self-compatible and undetermined self-incompatible. In addition if you have information about the genetic mechanism of self incompatibility, that is phenotypic studies of pollen germination on stigmas, genetic crosses, and/or identification of the genes involved in self-incompatibility that make possible the distinction between gametophytic and sporophytic incompatibility, score the type of incompatibility, then score gametophytic self-incompatible or sporophytic self-incompatible. Note that this only applies to genetic self-(in)compatibility. For instance, heterostylous species should not automatically be scored self-incompatible, nor dioecious species.

##### 84. Outcrossing rate (C1)

Value between 0 and 1. Note that outcrossing rate ¿ 1 can sometimes be given (due to statistical issues). In that case note 1. If the selfing rate is reported take 1 – selfing rate. If several populations have been measured in the same study then score all values (as separate entries). The additional table "Quality score" must be filled (not optional) with the method used to estimate outcrossing rate (corresponding to increasing quality)

1. Fis method, basically, outcrossing rate = (1 - Fis)/(1+Fis)
2. Population-based method: RMES (David et al., 2007) or INSTRUCT (Gao et al., 2007)
3. Pedigree-based method: Ritland's method = MLTR (Ritland, 2002) or BORICE (Koelling et al., 2012)

For pedigree-based methods, report the multi-locus outcrossing rate, not the single locus estimate (except if only one locus is used!). Additional equivalent methods exist but they are very rarely used. In addition the number type of genetic markers can be noted in the information table (recommended but not mandatory): allozyme, RAPD, RFLP, microsatellites, RadSeq, SNPs. When estimates are given for several populations, enter one value for each population. When Fis is used to infer outcrossing rate, make sure that individuals have been sampled in a single natural population. Don't consider collections of individuals (frequently noted as accessions) coming from a same "group" but not necessarily a population..

##### 90. Pollination syndrome (D1)

Inferring how flowers are effectively pollinated requires detailed studies, as many flower visitors might not contribute to pollination. Alternatively, pollination can be assumed from inflorescence or floral traits. Do not infer the syndrome yourself, but only used published descriptions. Please use the confidence level to indicate how pollinators were inferred :

1. (I have doubts about this record): inferred from flower/inflorescence traits
2. (I think this is right): observations of flower visitors
3. (I am certain this is right): pollination effectively demonstrated

Multiple pollinators are possible. For biotic pollination, we do not strictly follow phylogenetic categories for biotic pollinators, but rather intuitive categories, which might be paraphyletic. Possible pollination syndromes are:

**autonomous pollination** occurs when pollen is transferred to stigmas without other vectors. Examples are pollen falling on stigmas or direct contact between anthers and stigmas. Cleistogamous flowers necessarily fall in this category

**wind**

**water** (including rain)

**bee:** Hymenoptera clade Anthophila, which includes honey bees, bumblebees, and solitary bees

**beetle:** Coleoptera

**butterfly:** day-active lepidopterans

**moth:** night-active lepidopterans

**fly:** Diptera suborder Brachycera (excluding mosquitos).

**carrion fly and/or flesh fly:** Diptera families Calliphoridae and Sarcophagidae. These are flies whose larvae live in rotting meat and dung. They are attracted by flowers that imitate (through scent, colour, and/or texture) these substances

**wasp:** Hymenoptera of the Apocrita suborder that are not ants or bees. Includes social and solitary wasps, hornets, parasitic wasps, fig wasps.

**insect (other):** use if the pollinating insects belong to none of the above categories

**insect (general):** use if insect pollination is indicated without further precision on the insect taxon.

**bird**

**bat**

**non-flying mammal**

**other biotic pollinator**

##### 95. Dispersal syndrome (D1 )

As the morphological descriptions do not always capture the fruit-associated structures that determine dispersal mode (e.g. *Fragaria*, *Ficus*), we also score the dispersal syndrome as inferred from actual dispersal observations, or interpreted from the fruit characters. The trait states are copied from Perez-Harguindeguy et al. (2013):

**unassisted dispersal:** the seed or fruit has no obvious aids for longer-distance transport and merely falls passively from the plant.

**wind dispersal (anemochory)** includes (A) minute dust-like seeds (e.g. *Pyrola*, *Orchidaceae*), (B) seeds with pappus or other long hairs (e.g. willows (*Salix*), poplars (*Populus*), many *Asteraceae*), 'balloons' or comas (trichomes at the end of a seed), (C) flattened fruits or seeds with large 'wings', as seen in many shrubs and trees (e.g. *Acer*, birch (*Betula*), ash (*Fraxinus*), lime (*Tilia*), elm (*Ulmus*), pine (*Pinus*)); spores of ferns and related vascular cryptogams (Pteridophyta) and (D) 'tumbleweeds', where the whole plant or infructescence with ripe seeds is rolled over the ground by wind force, thereby distributing the seeds. The latter strategy is known from arid regions, e.g. *Baptisia lanceolata* in the south-eastern USA and *Anastatica hierochuntica* (rose-of-Jericho) in northern Africa and the Middle East.

**internal animal transport (endo-zoochory),** e.g. by birds, mammals, bats; many fleshy, often brightly coloured berries, arillate seeds, drupes and big fruits (often brightly coloured), that are evidently eaten by vertebrates and pass through the gut before the seeds enter the soil elsewhere (e.g. holly (*Ilex*), apple (*Malus*)).

**external animal transport (exo-zoochory);** fruits or seeds that become attached e.g. to animal hairs, feathers, legs, bills, aided by appendages such as hooks, barbs, awns, burs or sticky substances (e.g. burdock (*Arctium*), many grasses).

**dispersal by hoarding;** brown or green seeds or nuts that are hoarded and buried by mammals or birds. Tough, thick-walled, indehiscent nuts tend to be hoarded by mammals (e.g. hazelnuts (*Corylus*) by squirrels) and rounded, wingless seeds or nuts by birds (e.g. acorns (*Quercus spp.*) by jays).

**ant dispersal** (myrmecochory); dispersules with elaiosomes (specialised nutritious appendages) that make them attractive for capture, transport and use by ants or related insects.

**dispersal by water (hydrochory);** dispersules are adapted to prolonged floating on the water surface, aided for instance by corky tissues and low specific gravity (e.g. coconut).

**dispersal by launching (ballistichory);** restrained seeds that are launched away from the plant by 'explosion' as soon as the seed capsule opens (e.g. *Impatiens*).

**bristle contraction;** hygroscopic bristles on the dispersule that promote movement with varying humidity.

**no seeds or fruits:** to be used for species that are fully clonal and don't have dispersules

#### Flower-level characters

##### 100. Floral structural sex (D1)

From Schönenberger et al. (2020): "Flowers can be either bisexual (hermaphrodite) or unisexual. In our original study (Sauquet et al., 2017), we used a single character to distinguish among the many possible ways to be unisexual, depending on whether sterile organs of the opposite sex (staminodes or carpellodes) are found in flowers of a given sex, whether male and female flowers are found on the same plant (monoecy) or separate plants (dioecy), as well as the various intermediate combinations that exist (e.g., androdioecy, gynomonoecy). We have now simplified and divided this character into two and here capture only floral sex, distinguishing among three states: bisexual, incompletely unisexual (i.e., with pistillode in male flowers and/or staminodes in female flowers), and unisexual. All data from our previous dataset have now been rescored permanently into one of these three states. A second new character, Plant sexual system, resulting from the simplification outlined above, is excluded here because fossil flowers are usually found as individual, dispersed specimens, making it impossible to distinguish among different sexual systems such as monoecy and dioecy."

Added note: incompletely unisexual plant systems such as gynodioecy and androdioecy will typically be scored as polymorphic for floral structural sex. For instance, gynodioecious species with female and bisexual individuals (e.g., many *Dianthus* species, *Thymus*), each with structurally female and bisexual flowers, should be scored as unisexual / bisexual for this character. However, note that this character cannot be directly scored from plant sexual system without more information on floral structure, as gynodioecy and androdioecy can also refer to species with both unisexual and monoecious individuals. For instance, species of *Ficus* referred to as gynodioecious in the literature typically only have structurally unisexual flowers (with female individuals and figs and structurally bisexual but functionally male individuals and figs) and should be scored here as unisexual only. Likewise, plant sexual systems reported in the literature should be interpreted with caution, depending on whether structure or function is intended. For instance, *Amborella* is sometimes described as androdioecious, but is in fact functionally dioecious, having male individuals with structurally male flowers and female individuals with female flowers that are incompletely unisexual in structure (because they have 1-2 staminodes), but entirely unisexual in function. *Amborella* should be scored here as unisexual / incompletely unisexual.

##### 102. Ovary position (D1)

From Schönenberger et al. (2020): "The ovary is the part of the gynoecium where the ovules are produced. The ovary may be located on the receptacle and thus be positioned above the insertion level of the remaining floral organs (i.e., the ovary is superior and the flower is hypogynous). Alternatively, the ovary may be embedded in the receptacle and therefore be located below the insertion level of the remaining floral organs (i.e., the ovary is inferior and the flower is epigynous). Flowers with a hypanthium may either have a superior ovary (perigyny; e.g., many *Rosaceae*) or an inferior ovary (epiperigyny) (Simpson, 2010). It is also possible that the ovary is inferior to a certain degree only, such as half-inferior, if the receptacle is surrounding the ovary to its mid-level. Here we recorded the ovary position either as superior, inferior, or one of the following intermediate states: ¼ inferior or less, half-inferior, ¾ inferior or more."

##### 103. Flower length (C1)

In cm. Should be measured at anthesis, starting from the base of the receptacle or the base of an inferior ovary, excluding the pedicel.

##### 104. Flower diameter (C1)

In cm; to be measured at anthesis.

##### 107. Floral reward (D1)

The reward might be offered in the flowers or the inflorescences. The reward can be food for the visiting insects, such as pollen (including non-fertile pollen that is produced for reward purposes only), nectar, other sugar-containing liquids, oil (liquid fats), resin (solid or highly viscous secretions), food bodies (solid), or other tissues associated to the flower (e.g. stamens, bracts). Alternatively, food can be destined to larvae when the flowers serve as nurseries, either through dedicated structures (e.g. figs) or not: ovipositing insects of which the larvae feed on the ovules can nevertheless be pollinators. Alternatively, the reward offered by flowers can be shelter (e.g. sleeping places) or heat. Some flowers offer no reward, which is the case for abiotically pollinated flowers, but also for some attractive biotically pollinated, such as in the case of deceptive pollination. The precise mechanism of deceptive pollination can be described as a comment. See also Simpson and Neff (1981).

#### Perianth characters

##### 207. Symmetry of perianth (D1)

From Schönenberger et al. (2020): "There are many ways in which flowers can be zygomorphic (monosymmetric, with a single plane of bilateral symmetry). Here we record perianth symmetry, regardless of androecium or gynoecium symmetry; thus the character is not applicable when the perianth is absent. We distinguish strict actinomorphy (i.e., polysymmetry, with three or more planes of bilateral symmetry) from spiral actinomorphy. In addition, disymmetry (two orthogonal planes of bilateral symmetry; e.g., *Papaveraceae*) and asymmetry are treated here as separate character states. As for the fusion of the perianth, this character is applied to the perianth as a whole. In case of flowers with two or more perianth whorls, species were scored as actinomorphic if all whorls are actinomorphic and as zygomorphic if one or more whorls are zygomorphic."

##### 239. Main color of perianth at anthesis (D1)

This is the putative attractive color. If there is more than one color (either within the same organ or among organs), enter as separate character states. The choice is quite reduced compared to all variation in colors that exists, but the goal is to create broad categories for large-scale comparisons.

Yellow, green, cream, white, pink, brown, purple, red, grey, blue, orange. If the perianth is absent, this character should remain empty. If the most colourful organs are not the perianth, one should still indicate the colour of the perianth here, and indicate the identity and the colour of the attractive organs in traits 660 "Inflorescence attractive organ" and 661 "Inflorescence attractive colour". Note that a solitary flower is considered a one-flowered inflorescence.

#### Male reproductive function

##### 301. Number of fertile stamens (C1)

From Schönenberger et al. (2020): "Number of fertile (functional) stamens in bisexual or male flowers. Staminodes (co-occurring with fertile stamens) are not counted and female flowers are ignored for this character. Stamen number is highly variable within angiosperms and ranges from one (e.g., *Chloranthaceae*, Endress, 1987b) to several thousands (e.g., *Cactaceae*, Barthlott and Hunt, 1993). We record the number of fertile stamens in whorled or spiral flowers as a continuous character (with integer values of 1 and above). In cases of fusion among stamen whorls, the number of stamens may be difficult to determine. In such cases, additional information based on merism, anatomy, development, or comparison with closely related taxa may be taken into account. In synandria of Myristicaceae, for example, the number of fertile stamens can be deduced from the number of thecae present (Sauquet, 2003). In cases where stamen or anther morphology is not fully understood, we recorded the number of fertile stamens only when unequivocally clarified in the literature (e.g., *Malvaceae*, von Balthazar et al., 2004). Equivocal cases were left as missing data."

#### Female reproductive function

##### 401. Number of structural carpels (C1)

From Schönenberger et al. (2020): "Number of fertile or sterile carpels in bisexual or female flowers, recorded as a continuous character (with integer values of 1 and above). Contrary to the number of stamens (character 301), the number of co-occurring carpellodes (sterile carpels) is counted here because this number is often more easily obtained from the literature than the actual number of fertile carpels. However, consistently with our treatment of sexual dimorphism, the number of carpellodes in male flowers is ignored for this character. In multicarpellate, unilocular gynoecia with complete carpel fusion up to the stigma (e.g., *Primula*), it may be difficult to assess the number of carpels unequivocally. In such cases, we have scored the number of carpels only if it is well established based on anatomical or developmental investigations. Similarly, in gynoecia where one or more carpels are reduced (e. g., in the pseudomonomerous gynoecia of some *Arecaceae*, Stauffer et al., 2002), the total number of structural carpels was only scored when unequivocally determined in the literature. Contrary to the perianth and the androecium, we do not have a separate character for gynoecium merism here. This is because gynoecia with two or more whorls are rare in angiosperms and gynoecium merism therefore usually equals the number of carpels per flower."

##### 403. Fusion of ovaries (C1)

From Schönenberger et al. (2020): “Degree of ovary fusion expressed as a fraction of the total length of the ovary (from the floral base to the apex of the ovary). Fusion of styles and stigmas is not taken into account here. Not applicable when there is a single carpel.”

##### 411. Number of ovules per functional carpel (C1)

From Schönenberger et al. (2020): “Number of ovules per carpel recorded as a continuous character (with integer values of 1 and above). Reduced (sterile) carpels are not taken into account here.”

#### Inflorescences

##### 660. Inflorescence attractive organ (D1)

The attractive function can be fulfilled by the perianth or other organs. These can be bracts, leaves, stamens, or other organs (indicate in a comment), multiple organs (please leave a comment) or none. Note that single flowers should be considered as one-flowered inflorescences.

##### 661. Inflorescence attractive color (D1)

In many cases, the attractive function of the inflorescence is achieved by the perianth; thus, the attractive color is the same as the one scored in “239. Main color of perianth at anthesis”, and can be ignored here. In other cases, indicate the color of the bracts, stamens, etc. as scored in “660. Inflorescence attractive organ”. Single flowers can be considered as one-flowered inflorescences. If different colours exist, indicate either the dominant colour (if applicable) or each colour separately.

Note: for literature cited in Schönenberger et al. (2020), see that publication for full references.
